# Human NLRP1 is activated by ZAKɑ-driven ribotoxic stress response

**DOI:** 10.1101/2022.01.24.477516

**Authors:** Kim S. Robinson, Gee Ann Toh, Pritisha Rozario, Shima Bayat, Zijin Sun, Stefan Bauernfried, Rhea Nadkarni, Cassandra R. Harapas, Chrissie K. Lim, Werncui Chu, Kiat Yi Tan, Carine Bonnard, Radoslaw Sobota, John E. Connolly, Seth L. Masters, Kaiwen W. Chen, Lena Ho, Veit Hornung, Franklin L. Zhong

## Abstract

Human NLRP1 is a multifunctional inflammasome sensor predominantly expressed in skin and airway epithelium; however its function in skin-specific immunity and its mechanisms of activation are not fully understood. Here we report that human NLRP1 is specifically activated by the ZAKɑ- driven ribotoxic stress response pathway (RSR) induced by ultraviolet B (UVB) irradiation or select microbial ribotoxins. Biochemically, RSR-triggered NLRP1 activation requires ZAKɑ- dependent hyperphosphorylation of a human-specific linker region of NLRP1 (NLRP1^DR^), leading to the ‘functional degradation’ of the auto-inhibitory NLRP1 N-terminal fragment. Additionally, we show that fusing NLRP1^DR^ to the signaling domains of CARD8, which in itself is insensitive to RSR, creates a minimal inflammasome sensor for UVB and ribotoxins. In summary, these discoveries resolve the mechanisms of UVB sensing by human NLRP1, identify ZAKɑ-activating toxins as novel human NLRP1 activators, and establish NLRP1 inflammasome-dependent pyroptosis as an integral component of the ribotoxic stress response in primary human cells.

1. UVB-induced NLRP1 activation in human keratinocytes involves a nuclear DNA-independent stress response involving photodamaged RNA
2. ZAKɑ kinase is required for UVB-triggered, but not VbP- or dsRNA-induced human NLRP1 activation
3. ZAKɑ-activating microbial ribotoxins specifically activate the NLRP1 inflammasome in multiple primary human cell types
4. Hyperphosphorylation of a linker region (NLRP1^DR^) is required for RSR-dependent human NLRP1 activation

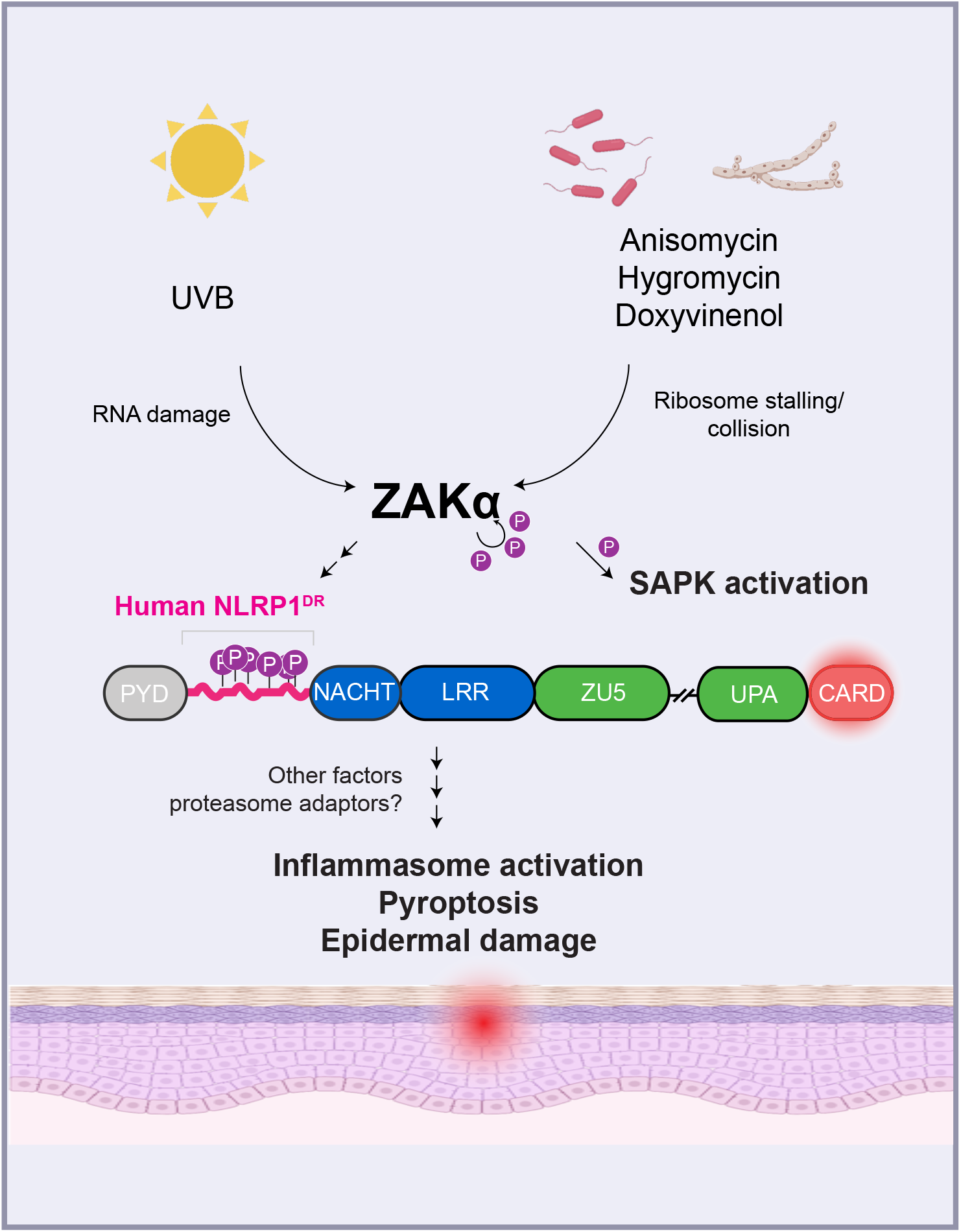

## INTRODUCTION

The innate immune system uses germline-encoded sensor proteins to recognize conserved pathogen- or damage-associated molecular patterns (PAMPs and DAMPs) (*1, 2*). One key innate immune system in vertebrates employs NACHT, LRR, and PYD domain-containing proteins (NLRPs) to sense and respond to pathogens and damage, particularly those that have gained access to the cytosol (*3–6*). Activated NLRPs then assemble the ‘inflammasome complex’, causing an inflammatory form of cell death known as pyroptosis, which is characterized by caspase-1 activation, GSDMD pore formation and IL-1 secretion (*3–10*). Human NLRP1 is notable among mammalian NLR sensors due to its unusual domain arrangement, high expression in non-hematopoietic organs and significant divergence from rodent counterparts (*11–13*). Although both human and mouse NLRP1 assemble the inflammasome through a conserved C-terminal CARD domain, they have evolved distinct ligand specificity. Chemical inhibitors of proteases DPP8 and DPP9, such as Val-boro-Pro (VbP) (*14, 15*) are the only known molecules that can activate both rodent and human NLRP1, in addition to a related inflammasome sensor CARD8 which shares a similar domain arrangement as NLRP1 (*16–18*). Our groups and others recently discovered that human NLRP1 senses enteroviral 3C proteases and double-stranded viral RNA, pointing to its role in antiviral immunity (*19–21*). Work from Beer and colleagues also showed that human NLRP1 detects ultraviolet B irradiation in skin keratinocytes (*22, 23*). None of these triggers activate rodent NLRP1s, which in turn senses bacterial and protozoan molecules, such as anthrax lethal factor (LF), *Shigella flexneri* IpaH7.8 and unknown molecule(s) from *Toxoplasma gondii* (*24–28*). Regardless of the species of origin, most known NLRP1 triggers require a common intermediate step to achieve full NLRP1/CARD8 inflammasome activation, namely the ‘functional degradation’ of the auto-inhibitory N-terminal fragment (*19, 20, 29–31*). This liberates the C-terminal UPA-CARD fragment to initiate inflammasome assembly (*17, 18, 32, 33*).

In contrast to other inflammasome sensors, human NLRP1 is predominantly expressed in the skin and airway epithelia (*19, 23, 34*). Rare germline mutations in NLRP1 cause Mendelian disorders with characteristic skin manifestations, with only the most severe cases demonstrating periodic fever and systemic auto-inflammation (*34–36*). In addition, a number of common NLRP1 SNPs confer increased risk for auto-immune skin disorders such as psoriasis (*37*). Thus human NLRP1 is likely to play a unique and specific role in skin immunity that is not shared by other inflammasome sensors. In contrast to humans, several studies have established murine skin epidermis expresses either very low or undetectable levels of NLRP1 inflammasome components (*23, 38*). Thus, the role of NLRP1 inflammasome in skin is not conserved in mice. Amongst all human NLRP1 triggers identified thus far and supported by genetic evidence, ultraviolet (UV) irradiation with wavelengths 280-315 nm, or UVB is the most relevant to human skin. Importantly, work from the Beer group and others have established that UVB-dependent inflammasome activation in human keratinocytes is completely dependent on NLRP1 and does not involve NLRP3 (*22, 23*). Furthermore, UVB does not activate NLRP1 inflammasome activation in murine keratinocytes (*23*). However, the molecular mechanisms by which human NLRP1 senses UVB have not been explored in detail. In particular, it is not clear if NLRP1 senses UVB irradiation directly, or responds to a cellular danger signal or signaling event that is induced by UVB.

Ultraviolet solar irradiation is divided into three wavelength bands: UVC at 100 −280 nm, UVB at 280-315 nm and UVA at 315-400 nm (*39*). UVB is responsible for most of the acute effects of sunburn and predominantly affects the epidermal layer of human skin consisting of keratinocytes (Figure 1A). At the cellular level, the principal signalling response triggered by UVB is thought to be initiated in the nucleus, where UVB causes DNA photo-adducts and DNA breaks (*39–42*). However, earlier work pointed to the existence of other UVB response pathways that operate in parallel to, and independently of DNA damage and repair. For instance, in human cancer cell lines, UVB causes rapid stress-activation protein kinases (SAPK) activation and the transcription of pro-inflammatory genes (*43–48*). This reaction could be recapitulated in enucleated cells and cytoplasmic extract, suggesting that it does not originate from nuclear DNA damage (*49*). Although the full molecular details of this cytosolic UVB pathway await further investigation, several studies in the literature implicate RNA damage and the translational machinery as key intermediates. Of note, the action spectra of UV (measured by its effects on gene activation, cytotoxicity and carcinogenesis) coincide with the absorption maxima of not only DNA, but also RNA (*43, 50*). UVB has been shown to cause damage to the 28s ribosomal RNA, leading to ribosome stalling and/or collisions (*51, 52*). It is now known that ribosome stalling/collisions can activate at least three distinct, genetically defined stress pathways, namely Ribosome Quality Control (RQC), the Integrated Stress Response (ISR) and Ribotoxic Stress Response (RSR) (*53*– *57*). Operating together, these pathways monitor the ‘health’ of the translational machinery. Once activated, they can either ensure cell survival by repairing damaged ribosomes or trigger cell death (*58, 59*). More recently, the long splice isoform of the MAP3K20 gene product, ZAKɑ was discovered to be the proximal sensor of the RSR in human cells (*56, 57*). Most importantly, ZAKɑ is genetically required for UV-triggered SAPK activation. ZAKɑ-dependent SAPK activation could even be recapitulated by cytosolic delivery of UVC-damaged heterologous RNA, suggesting that this response is truly DNA independent, cell-intrinsic (*57*), and is distinct from the known mode of detection of damaged noncoding RNA by TLR3, which operates in a non-cell-autonomous manner (*60, 61*). These seminal findings by the Green and Bekker-Jensen labs establish ZAKɑ as a critical regulator of UVB induced cell-intrinsic stress response. In this study, we build upon this knowledge and report an unexpected connection between the ZAKɑ-dependent RSR and NLRP1 inflammasome-driven pyroptosis. We demonstrate that ZAKɑ is indispensable for UVB sensing by the human NLRP1 inflammasome, and RSR-inducing agents, such as certain microbial ribotoxins, are selective NLRP1 inflammasome agonists in multiple primary human cell types. Mechanistically, UVB and other ZAKɑ-activating agents activate human NLRP1 via the hyperphosphorylation of a species-specific linker region, which is dispensable for VbP-triggered NLRP1 activation. Our results thus resolve the mechanism of UVB sensing by the human NLRP1 inflammasome in skin epidermis, and identify a new general pathway of NLRP1 activation by ZAKɑ-activating agents.

**Figure 1.**
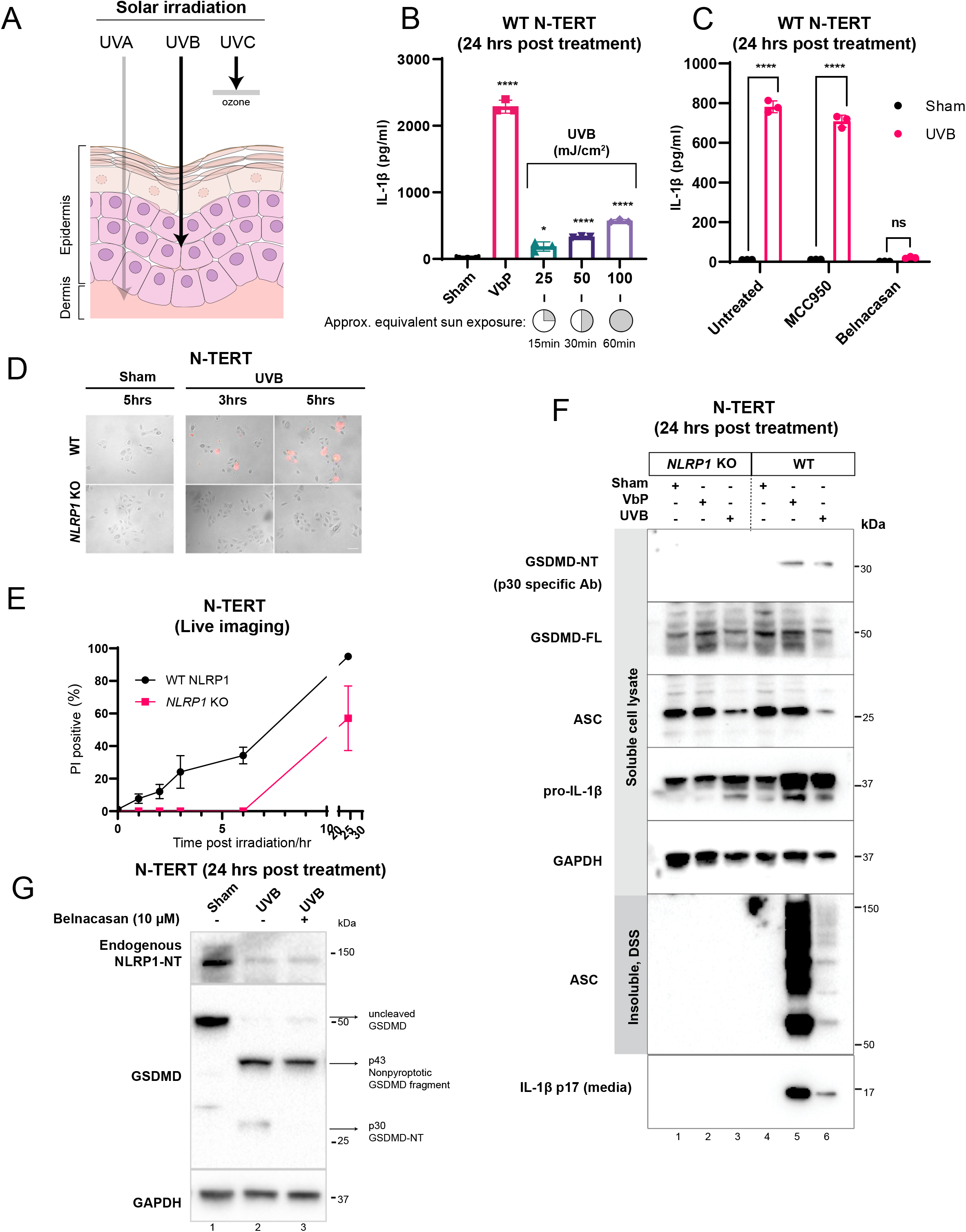
Validation of UVB as a specific trigger for the human NLRP1 inflammasome. A. Schematic showing depth penetration of solar irradiation (composed of UVA, UVB and UVC) into human skin (composed of epidermis at the top and dermis at the bottom). B. IL-1β ELISA of N-TERT cell media collected 24 hours after treatment with: Sham (control for heat generated during UVB exposure), VbP (3 μM) or increasing doses of UVB (25, 50, 100 mJ/cm^2^). The approximate equivalent sun exposure times of these doses is indicated as referenced (*62*). C. IL-1β ELISA of N-TERT cell media collected 24 hours after being either sham or UVB (100 mJ/cm^2^) irradiated. Cells were pre-treated with MCC950 (5μM) (2 hours) or Belnacasan (10 μM) (30 mins). D. Representative images of PI positive wild-type or *NLRP1* KO N-TERT cells after UVB (100 mJ/cm^2^) irradiation. Cells were grown in 0.25 μg/mL PI. Representative images taken at 3 or 5 hours following UVB irradiation, and 5 hours sham irradiation. E. Quantification of percentage of PI positive wild type N-TERT cells or *NLRP1* KO N-TERT cells shown in D. Cells imaged after 2,3,4,5,6 and 24 hours after UVB irradiation. F. Immunoblot of wild type N-TERT cells or *NLRP1* KO N-TERT cells treated with VbP (3 μM), UVB (100 mJ/cm^2^) or sham irradiation. Note that the GSDMD-NT Ab is specific for the p30 fragment and does not recognize other cleaved products such as p43 or p20. G. Immunoblot of wild type N-TERT cells or *NLRP1* KO N-TERT cells treated with UVB (100 mJ/cm^2^) or sham irradiated. For lane 3, cells were treated with Belnacasan (10μM) for 30 mins before UVB irradiation. The GSDMD Ab used here detects all forms of GSDMD, including full length, p43 and p30.

## RESULTS

### Validation of UVB as a specific trigger for the NLRP1 inflammasome

Using N-TERT immortalized human keratinocytes (hereby referred to as N-TERTs), we confirmed published findings that physiologically relevant levels of UVB irradiation (25-100 mJ/cm^2^) caused IL-1β secretion in a dose-dependent manner (Figure 1B) (*22*). At 100 mJ/cm^2^ (equivalent to approximately 60 mins of direct midday European summer sun exposure) (*62*), UVB caused approximately 30% of IL-1β secretion relative to a near saturating dose of Val-boro-Pro (VbP) (3 μM), a specific agonist for the NLRP1 inflammasome (Figure 1B). No significant IL-1β secretion was observed in the ‘sham’ irradiated sample, which was used to control for the heat generated by the UV lamp. UVB-induced IL-1β secretion could be blocked by caspase-1 inhibitor, Belnacasan (10 μM) but not NLRP3 inhibitor MCC950 (3 μM) (Figure 1C). Of note, *NLRP3* mRNA is undetectable in N-TERT cells and primary human keratinocytes (*19*). In agreement with Fenini et al., (2018), CRISPR/Cas9 deletion of the *NLRP1* gene completely abrogated UVB-induced IL-1β secretion (Figure S1B), which was fully restored by stable NLRP1 overexpression in the KO background (Figure S1B) (*22*). Furthermore, in wild-type N-TERT cells, UVB elicited rapid propidium iodide (PI) uptake (within 3 hours) (Figure 1D, 1E), which occurred even more rapidly than the established NLRP1 agonist VbP (∼6 hours in wild-type N-TERT cells, see additional evidence in Figure 4-7) (*20*). The PI uptake at earlier time points (3 hours) was abrogated in *NLRP1* KO cells, which incorporated PI with much slower kinetics (Figure 1D, 1E). Hence we used PI uptake at 3 hours as one of the readouts to distinguish pyroptosis from other forms of cell death in N-TERT cells.

Next, we examined the biochemical hallmarks of inflammasome activation in UVB-treated N-TERT cells. Similar to VbP, UVB caused accumulation of detergent insoluble ASC oligomers and GSDMD p30 cleavage in wild-type, but not *NLRP1* KO cells (Figure 1F, lanes 5 and 6 vs. lanes 2 and 3). Using a p17 cleavage specific antibody, we validated that UVB indeed caused the secretion of the bioactive, p17 form of IL-1β, i.e. the specific caspase-1 cleavage product, in an NLRP1-dependent manner (Figure 1F, lanes 5 and 6 vs. 2 and 3, bottom panel). Taken together with published observations, these results confirm that UVB indeed functions as a specific NLRP1 trigger in human keratinocytes. Our data do not suggest that *NLRP1* KO or inflammasome inhibition could prevent all the cellular effects of UVB. For instance, *NLRP1* KO cells still slowly incorporated PI after UVB exposure (Figure 1E, S1A), which is consistent with late stage apoptosis/necrosis. Rather, we propose that NLRP1 inflammasome activation is one of the major consequences of UVB irradiation in human skin keratinocytes.

As further support for specific NLRP1 activation by UVB, we found that the level of the endogenous NLRP1 N-terminal fragment (NLRP1-NT) decreased significantly after UVB exposure (Figure 1G, lane 1 vs. 2). This occurred without any significant change in *NLRP1* mRNA level (mean RNAseq FKPM 7.6 vs 6.8 before and after UVB, p>0.05) and could not be prevented by Belnacasan (10 μM), which selectively abrogated caspase-1-dependent GSDMD p30 cleavage, but not the non-pyroptotic p43 cleavage (Figure 1G, lane 2 vs. 3). These results suggest that NLRP1-NT was most likely degraded in a post-translational step upstream of caspase-1 activation. Thus, just like all other established NLRP1 agonists such as VbP, dsRNA and enteroviral 3C protease, UVB likely leads to the ‘functional degradation’ of NLRP1-NT.

### UVB-dependent NLRP1 activation likely involves a nuclear DNA-independent stress response caused by photodamaged RNA

NLRP1 itself is unlikely to function as a direct UVB sensor, as the structure of the NLRP1:DPP9 complex, recently reported by us and others, does not harbor any features of light sensing proteins (*17, 18*). Hence, UVB-triggered NLRP1 activation likely requires additional cellular factors. To elucidate this potentially novel pathway, we devised two approaches: 1) to narrow down and identify the exact identity of the UVB-dependent ‘danger molecules’ that trigger NLRP1 and 2) to genetically define the obligate regulator(s) that relay the signal from such molecules to NLRP1. For the former, we considered three major types of UVB-elicited cellular damage: DNA breaks, RNA photodamage and oxidative damage by free radicals (which are not necessarily mutually exclusive) (Figure 2A). UVB is one of the most studied DNA mutagens and induces DNA photo-adducts such as cyclobutane pyrimidine dimers (CPDs) and pyrimidine (6–4) pyrimidone photoproducts (64PPs). These lesions are sensed and repaired by dedicated pathways such as nucleotide excision repair (NER). Defects in NER result in over-accumulation of ssDNA and dsDNA breaks and cause human Mendelian photosensitivity disorders such as Xeroderma Pigmentosum (XP) and Cockayne syndrome (CS), as well as common forms of skin cancer. We therefore tested if DNA damage itself was sufficient to explain UVB-induced NLRP1 activation by treating wild-type and NLRP1 KO N-TERT cells with DNA damaging chemicals, camptothecin (CPT), etoposide and cisplatin (Figure 2B). These drugs were chosen as they could induce different types of DNA lesions; in particular, cisplatin induces intra-strand adducts which, just like UVB photo-adducts, are repaired by NER. Over 24 hours of treatment, none of these chemicals induced IL-1β secretion, except etoposide at high concentrations (>10 μM). However, this was not eliminated, but in fact increased in NLRP1 KO cells. In a similar experiment, H_2_O_2_, a free radical inducer, also failed to cause significant IL-1β secretion in N-TERTs, despite a high level of cytotoxicity over 24 hours. In addition, none of the chemicals tested demonstrated differential kinetics of PI uptake between wild-type and NLRP1 KO cells (Figure S2A). These results suggest that DNA damage, or oxidative damage by free radicals alone cannot account for UVB-induced NLRP1 inflammasome activation in human keratinocytes, although we could not rule out that they act as modifiers of this process. We next considered the possibility that unrepaired DNA photo-adducts themselves act as intracellular ligands for NLRP1. To test this, we deleted the *DDB2* gene in N-TERT cells (Figure S2C). DDB2 is the proximal sensor for UVB-elicited DNA lesions and initiates NER (Figure S2B). Its deletion inactivates the NER pathway and leads to accumulation of unrepaired photo-adducts, as seen in patients with Xeroderma pigmentosum-E caused by germline DDB2 loss-of-function mutations. As compared to wild-type cells, *DDB2*-null N-TERTs did not show any significant difference in the amount of IL-1β secretion following UVB despite accelerated PI uptake and cell death (Figure S2D, S2E). Therefore, UVB-induced DNA damage or the subsequent NER pathway are not the primary cause for NLRP1 inflammasome activation.

**Figure 2.**
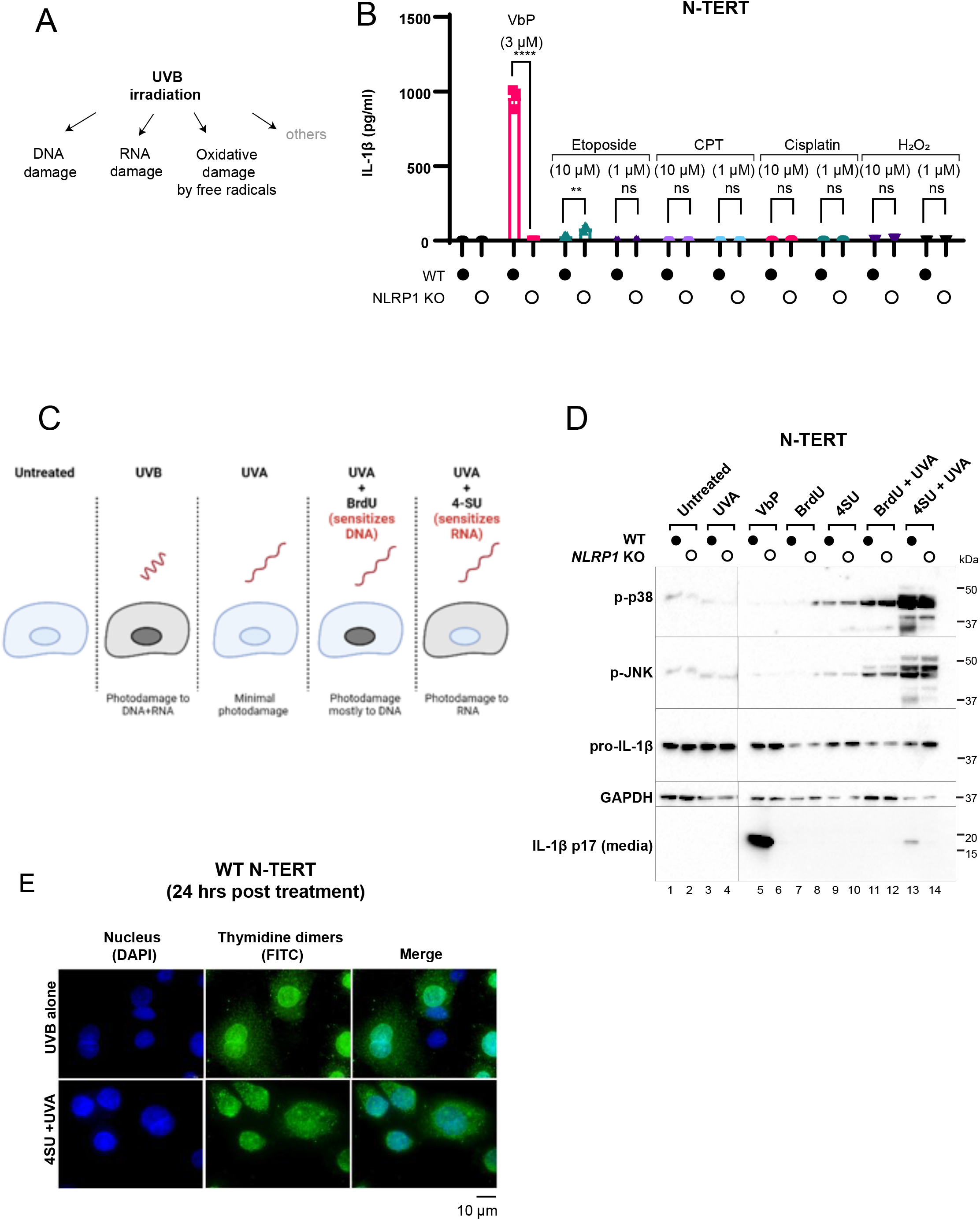
UVB-dependent NLRP1 activation likely involves a nuclear DNA-independent stress response induced by photodamaged RNA. A. Schematic indicating types of cellular damage caused by UVB irradiation. B. IL-1β ELISA of N-TERT cell media collected 24 hours after being treated with indicated drugs. C. Experimental approach to selectively sensitize either DNA and RNA to UV using nucleoside/nucleotide analogs. D. Immunoblot of wild type N-TERT cells or *NLRP1* KO N-TERT cells treated with the indicated combinations of photosensitizer and UV. VbP (3 μM) was included as a positive control. E. Immunofluorescence staining of N-TERT cells treated with either UVB (100mJ/cm^2^) or 4-SU (1 μM) + UVA (500mJ/cm^2^). Cells were fixed immediately post irradiation and stained for UV DNA damage with an anti-thymidine dimer antibody (green) and counterstained with DAPI (blue).

Next we considered another source of UVB induced cellular damage: RNA photodamage. Similar to DNA, RNA is also excited by UVB. However, the physiological consequences of RNA photodamage in human cells are much less understood. To preferentially elicit RNA photodamage over DNA damage, we employed a selective sensitization technique first developed to study RNA-protein interactions (e.g. PAR-CLIP (*63*)) (Figure 2C). Nucleoside analog 4-thiouridine (4-SU) incorporates selectively into cellular RNAs (but not DNA) and allows the substituted RNA species to be excited by UVA, which does not affect unmodified nucleic acids due to its lower energy per photon. Conversely, 5-bromo-2’-deoxyuridine (BrdU), a thymidine analog, selectively sensitizes DNA (but not RNA) to UVA irradiation. Using thymidine dimer immunostaining as a marker for DNA damage, we first validated this approach by showing 4-SU+UVA did not significantly damage nuclear DNA (Figure 2E). Next we subjected wild-type and NLRP1 KO N-TERT cells to combinations of UVA with either 4-SU and BrdU. Of all the conditions tested, only 4-SU+UVA, which selectively damages RNA, recapitulated IL-1β p17 secretion 24 hours post-irradiation. Most importantly, this effect was abrogated by NLRP1 deletion (Figure 2D, lane 13 vs. lane 14, bottom panel). These results demonstrate that UV-dependent RNA photodamage is sufficient to activate NLRP1 in human skin keratinocytes. This unusual requirement for RNA was reminiscent of the previously reported UV-driven SAPK activation, which was also shown to involve only cytosolic factors (*49*). Indeed, 4-SU+UVA led to the most robust SAPK (p38 and JNK) phosphorylation among the conditions tested (Figure 2D, lanes 13 and 14 vs lanes 1-12, upper panels). Based on these considerations, we speculated that UVB-triggered NLRP1 inflammasome activation and SAPK activation share the common mechanism involving UV-damaged RNA. Thus we went on to genetically examine the precise upstream regulators for UVB-triggered NLRP1 activation in subsequent experiments.

### RSR kinase ZAKɑ selectively controls UVB-dependently NLRP1 activation

Recently, the proximal sensor for UVB-triggered SAPK activation was found to be the long splice isoform of the *MAP3K20* gene product, also known as ZAKɑ. ZAKɑ is not known to sense nucleic lesions directly, but rather senses aberrant ribosomes that have stalled and/or collided after encountering a translocation-blocking mRNA lesion, such as those induced by UVB. Activated ZAKɑ undergoes extensive self-phosphorylation and phosphorylates downstream SAPKs such as p38 and JNK, which in turn trigger inflammatory signaling and/or cell death (Figure 3A) (*56, 57*). Collectively, this pathway was termed the ribotoxic stress response (RSR). Due to its shared involvement for RNA damage, we examined whether RSR intersects with UVB-induced NLRP1 activation.

**Figure 3.**
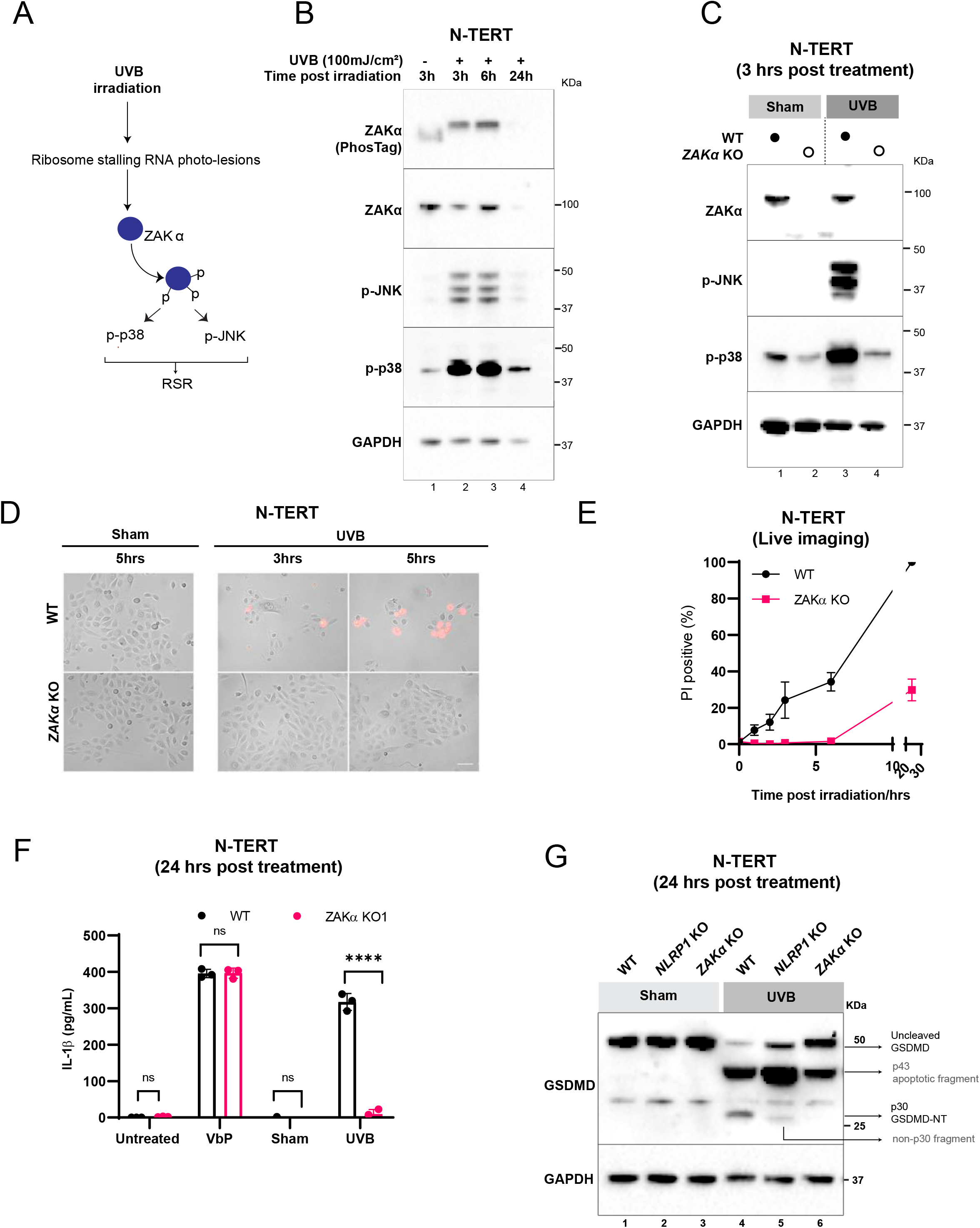
RSR kinase ZAKɑ controls UVB-but not VbP- or dsRNA-triggered NLRP1 activation. A. Schematic showing ribosomal stress response (RSR) signalling through ZAKɑ and its downstream effects on SAPKs, p38 and JNK. B. Immunoblot of N-TERT cells at the indicated time points after UVB (100mJ/cm^2^) irradiation. C. Immunoblot for ZAKɑ, p-p38 and p-JNK from WT or *ZAKɑ* KO N-TERT cells 3 hours after UVB (100mJ/cm^2^) irradiation. D. Images of PI positive wild type cells or *ZAKɑ* KO N-TERT cells after sham or UVB (100mJ/cm^2^) irradiation. E. Quantification of the percentage of PI positive wild type or *ZAKɑ* KO N-TERT cells following UVB (100mJ/cm^2^) irradiation. Cells imaged after 2,3,4,5,6 and 24 hours after UVB irradiation. F. IL-1β ELISA of WT or *ZAKɑ* KO N-TERT cell culture media 24 hours after sham or UVB (100mJ/cm^2^) irradiation. G. GSDMD Immunoblot of WT, *ZAKɑ* KO or *NLRP1* KO N-TERT cells treated with UVB (100mJ/cm^2^) or sham irradiated. Cell lysates were harvested 24 hours later. Different GSDMD cleavage fragments are shown by black arrows. Note that the GSDMD Ab used in this experiment recognizes all GSDMD cleaved products. In *NLRP1* KO cells UVB leads to a weak band <30kDa, which cannot be detected by p30 specific Ab shown in Figure 1F. Hence it should not be confused with the caspase-1 dependent p30 fragment.

We first showed that UVB indeed led to *bona fide* RSR activation within N-TERT keratinocytes. In contrast to certain commonly used cancer cell lines, N-TERT cells demonstrate basal levels of p38 phosphorylation. Regardless, UVB (100 mJ/cm^2^) led to prominent increase in phosphorylation of all detectable p38 and JNK isoforms, as well as hyperphosphorylation of ZAKɑ itself (Figure 3B, lanes 1 vs 2-3) within 3 hours. Vind and colleagues additionally reported that activated phosphorylated ZAKɑ (p-ZAKɑ) undergoes proteasomal degradation as the result of a *de novo* phospho-degron (*56*). This effect was especially pronounced in UVB-treated N-TERT cells, as total ZAKɑ was no longer detectable at 24 hours post-UVB exposure (Figure 3B, lanes 1 vs 4). In this context, we also validated that selective RNA photodamage by 4-SU+BrdU-UVA also triggered robust ZAKɑ activation. Notably, only N-TERT cells treated with 4-SU+UVA (which sensitizes RNA to photodamage), but not those treated with BrdU+UVA demonstrated ZAKɑ hyperphosphorylation and degradation (Figure S3A). Thus, the conditions that activate ZAKɑ-mediated RSR coincide with those required to activate the NLRP1 inflammasome in N-TERT cells.

We then hypothesized that ZAKɑ is upstream of NLRP1 activation by UVB. To test this, we generated *ZAKɑ* KO N-TERT cells (Figure S3B). In agreement with the original reports (*56, 57*), ZAKɑ deletion abrogated RSR signaling, as evidenced by the absence of p-p38 and p-JNK induction following UVB irradiation (Figure 3C, lanes 3 vs 4), but did not affect endogenous caspase-1 and pro-IL-1β expression (Figure S3B). Strikingly, *ZAKɑ*-null N-TERT cells no longer underwent inflammasome-driven pyroptosis after UVB treatment. This was supported by 1) the loss of IL-1β secretion (Figure 3F, S3C), 2) markedly delayed PI uptake (Figure 3D, 3E) and 3) the selective loss of GSDMD p30 cleavage in UVB-treated *ZAKɑ* KO cells (Figure 3G, lane 1 vs. 6). In addition, acute chemical inhibition of the ZAKɑ kinase activity using known inhibitor Nilotinib also abrogated IL-1β p17, mimicking *ZAKɑ* KO (Figure S3D). The effect of ZAKɑ on the NLRP1 inflammasome was restricted to UVB and did not extend to other NLRP1 activators, such as VbP and intracellular poly(I:C), as no difference in IL-1β secretion or GSDMD p30 cleavage was observed between wild-type and *ZAKɑ* KO N-TERT cells stimulated with these triggers (Figure 3F, S3E-F, S6A). Taken together, these results suggest that ZAKɑ is selectively required for the NLRP1 inflammasome activation downstream of UVB.

### Orthogonal means of ZAKɑ activation activates NLRP1 inflammasome in diverse primary human cell types

Next we carried out ‘sufficiency’ experiments to test if orthogonal methods of ZAKɑ activation other than UVB could also activate the NLRP1 inflammasome. We employed two independent systems of manipulating ZAKɑ that did not involve UVB. In the first, ZAKɑ and its truncation mutants constructed by Vind et al were overexpressed along with NLRP1 in a 293T-ASC-GFP reporter cell line (Figure 4A, 4B, S4A). Although this system required overexpression of inflammasome components, it offered sufficient dynamic range and sensitivity to distinguish true NLRP1 activators from nonspecific cytotoxic compounds that could theoretically lead to artifactual ASC-GFP aggregation. In addition, as 293T cells do not express endogenous caspase-1 or GSDMD, this reporter allowed a more direct test for NLRP1 regulators, without the interference of ensuing pyroptosis. The same 293T-ASC-GFP cell line and similar ones had been used to identify VbP and 3Cpro as human NLRP1 activators, and also to validate the NLRP1-DPP9 complex structure (*14, 17–19, 32*). When co-expressed with wild-type NLRP1, full-length ZAKɑ led to marked increase in the number of cells displaying ASC-GFP specks (Figure 4C). This effect required the C-terminal sensor and ribosome binding domains (S and CTD, respectively), as well as an intact kinase domain (Figure 4A, 4C), which are all required for ZAKɑ to initiate RSR signaling (*56, 57*). The short splice isoform of *MAP3K20*, ZAKβ was unable to activate NLRP1-driven ASC-GFP speck formation. This result was particularly noteworthy because ZAKɑ and ZAKβ share an identical kinase domain (Figure S4B); however, ZAKɑ has a unique C-terminus that associates with the ribosome and participates in RSR (Figure S4C). A patient-derived mutation in the ZAKɑ-specific SAM domain (p.Phe368Cys), which was shown to be hyperactive in RSR initiation (*56, 64*), also activated NLRP1 in 293T-ASC-GFP cells (Figure 4C). These findings demonstrated in an orthogonal cell line that overexpressing ZAKɑ triggers NLRP1 activation. In addition, the ability of ZAKɑ to activate NLRP1 requires the same domains as those needed to initiate RSR.

**Figure 4.**
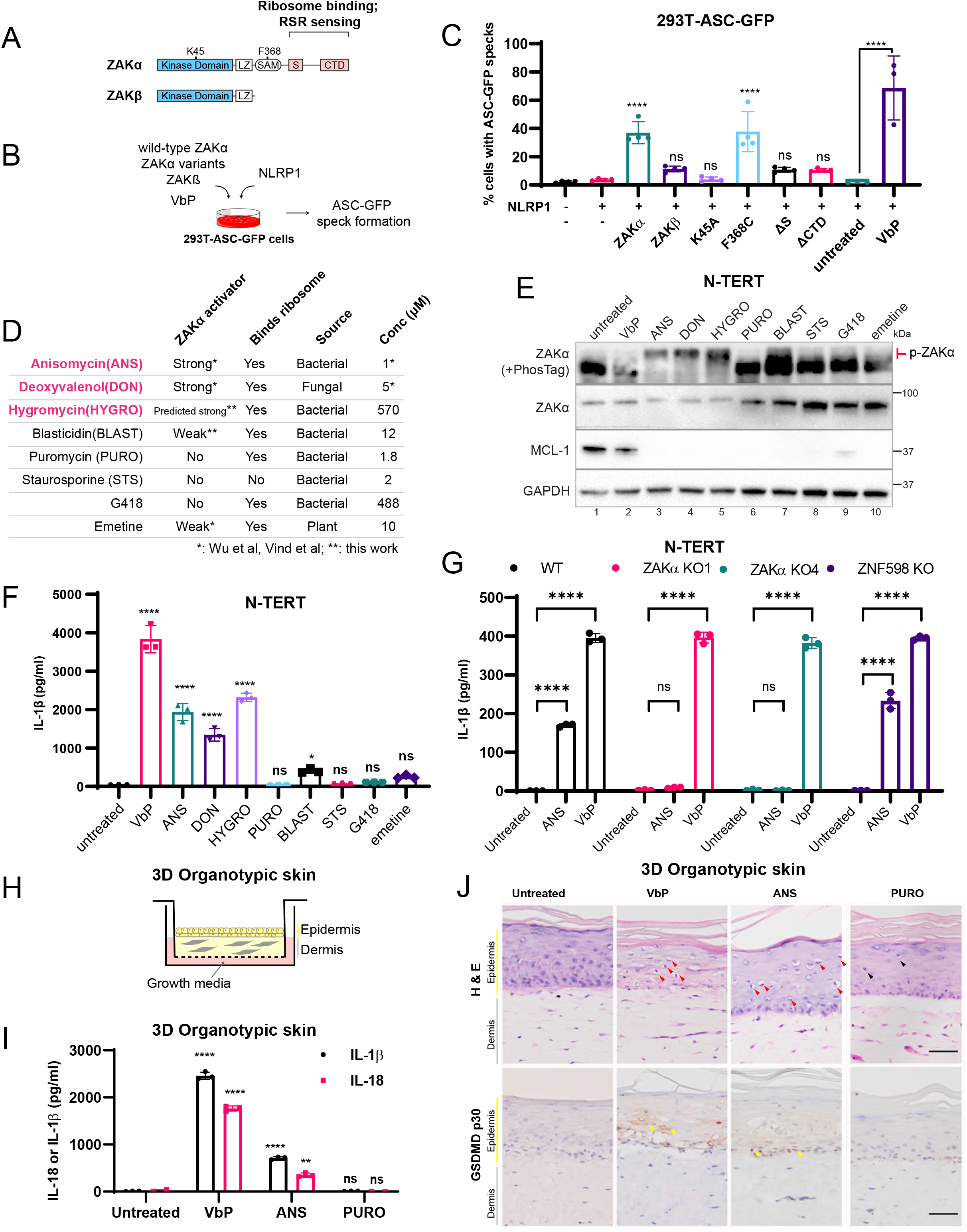
Orthogonal means of ZAKɑ activation triggers the NLRP1 inflammasome. A. Domain arrangements of ZAKɑ and ZAKβ, which result from alternatively spliced mRNAs from the MAP3K20 gene. ZAKɑ domains involved in RSR and ribosome binding are indicated. K45: ZAKɑ kinase catalytic residue; F368: residue mutated in split-hand- and-foot syndrome. B. Experimental setup to test the effect of ZAKɑ variants and ZAKβ on NLRP1 in 293T-ASC-GFP reporter cells. C. Quantification of the percentage of 293T-ASC-GFP cells showing ASC-GFP specks. Cells were transfected with the indicated constructs or treated with VbP, and fixed 24 hours post transfection/treatment. ASC-GFP specks were quantified from wide-field fluorescence microscopy images (20x magnification) with >200 cells per condition. D. List of ZAKɑ activating compounds and negative control cytotoxic drugs used in this study. The strengths of the known ZAKɑ activators, ANS, DON and emetine were adapted from Vind et al and Wu et al. The effects of HYGRO and BLAST on ZAKɑ were unknown before this study, to the best our knowledge. E. Immunoblot of N-TERT cell lysate after 3 hour treatment with the indicated drugs. The concentrations used are listed in D. Note that ZAKɑ phosphorylation can be detected by the subtle upshift on SDS-PAGE alone but becomes more obvious on PhosTag SDS-PAGE (marked in red). F. IL-1β ELISA of N-TERT cell media after 24 hours of treatment with the indicated drugs at concentrations specified in D. Note that a smaller volume of the media and a higher number of cells were used in this experiment, accounting for the overall higher concentration of IL-1β. G. IL-1β ELISA of culture media from N-TERT cells of the indicated genotypes after 24 hours of drug treatment. VbP was used at 3 μM. H. Graphical representation of 3D organotypic skin cultures consisting of stratified epidermis on collagen embedded dermis. I. IL-1β and IL-18 ELISA of culture media from 3D organotypic skin cultures treated with the indicated drugs. J. H&E and cleaved GSDMD-NT (p30 specific) immunostaining of 3D organotypic skin cultures treated with the indicated drugs. Scale bar = 100μm. Red arrows indicate dyskeratotic keratinocytes with diminished eosin but dense hematoxylin staining that were abundant in VbP- and ANS-treated cultures. Black arrows indicate putatively apoptotic cells with low eosin and hematoxylin staining that were abundant in PURO-treated samples. Yellow arrows indicate membranous GSDMD p30 staining.

In the second approach, we tested the potential effects of established ZAKɑ activating toxins, anisomycin (ANS) and doxyvinenol (DON) to activate the NLRP1 inflammasome. ANS and DON share a similar mechanism of action, i.e. by directly targeting the peptidyl transfer center (PTC) of elongating ribosomes, thus mimicking the effect of UVB-damaged RNA in causing ribosome stalling and subsequent collisions. To control for their inhibitory action on *de novo* protein synthesis, we compared ANS and DON to other well-established translational inhibitors (BLAST-blasticidin, PURO-puromycin, emetine and G418) which target different sites of the ribosome complex and therefore do *not* activate ZAKɑ. We also included a nonspecific cytotoxic drug (STS-staurosporine) to rule out the the possibility that NLRP1 could be indirectly activated by apoptosis, although we considered this unlikely, since none of the cytotoxic DNA damaging compounds activated NLRP1 (Figure 2B). We put our hypothesis to a more stringent test by including hygromycin (HYGRO), whose effect on ZAKɑ was unknown at the time of our study. Based on early work with yeast ribosomes (*65*), we predicted HYGRO to be a ZAKɑ-RSR activator by targeting an overlapping site on human ribosomes as ANS and DON. If ZAKɑ were truly a necessary and sufficient activator for NLRP1, only ANS, DON and HYGRO should specifically induce NLRP1-dependent pyroptosis in N-TERT cells. For most of the drugs, we chose concentrations characterized in detail in Vind et al., HYGRO and STS were used at concentrations widely chosen to elicit cytotoxicity after 16-24 hours of treatment (Figure 4D). We first confirmed that ANS, HYGRO and DON indeed led to ZAKɑ phosphorylation in N-TERT cells at 3 hours post treatment (Figure 4E, lanes 3, 4 and 5), whereas none of the other drugs could do so, with the possible exception of BLAST, although its effect was far less pronounced. By contrast, the level of anti-apoptotic protein MCL-1, a well established sensor for translational inhibition, was reduced by all drugs. This control experiment, in addition to those shown in Vind et al., and Wu et al., demonstrated that the ability of ANS, HYGRO and DON to activate ZAKɑ was unique and not caused by general, non-specific translation inhibition.

Out of the panel of drugs tested, only ANS, HYGRO and DON robustly activated the inflammasome-driven IL-1β secretion in N-TERT cells after a 24 hour incubation. BLAST led to a modest amount of IL-1β, in agreement with its role as a weak ZAKɑ activator. None of the non-ZAKɑ activating drugs caused IL-1β release (Figure 4F, S4E). In subsequent epistasis experiments, we confirmed that ANS-dependent inflammasome activation was completely dependent on ZAKɑ and NLRP1, as CRISPR/Cas9 deletion of either *ZAKɑ* or *NLRP1,* or chemical inhibition of ZAKɑ kinase activity by Nilotinib (0.1 μM), eliminated all hallmarks of inflammasome-driven pyroptosis including 1) IL-1β secretion 24 hours after treatment (Figure 4G), 2) rapid PI uptake within 3 hours (Figure S4H), 3) ASC polymerization (Figure S4G, lanes 2 vs 5) and 4) GSDMD cleavage (Figure S6B, lanes 6 vs 8). Re-expressing wild-type NLRP1 restored ANS-triggered ASC-oligomerization, IL-1β release and lytic cell death in NLRP1 KO cells (Figure S4H, S4I). Thus the ability to trigger NLRP1 inflammasome is unique to RSR-activating drugs (ANS, DON and HYGRO) and is mediated entirely by ZAKɑ. Global translational inhibitors, which do not activate ZAKɑ (Figure 4E, lanes 6-10), cannot activate the NLRP1 inflammasome in N-TERT cells. ANS also functions as a ZAKɑ-dependent NLRP1 activator in 293T cells, using ASC-GFP speck formation as well as NLRP1 CT (UPA-CARD) self-oligomerization as readouts (Figure S4J, S4K, S4M). Partial deletion of the ribosome quality control factor ZNF598 did not affect ANS- or VbP-triggered NLRP1 activation in N-TERT cells (Figure 4G, S4C-D). We also confirmed in a panel of KO N-TERT cells that the ability of ANS to activate the inflammasome was independent of *NLRP3* but required all effector components of the canonical inflammasome including ASC, CASP1 and GSDMD (Figure S4F). These results provide further evidence that ZAKɑ is a specific upstream regulator of the NLRP1 inflammasome in human keratinocytes.

To reiterate, the effect of ANS, DON and HYGRO was not caused by general translational inhibition, as non-ZAKɑ-activating translational inhibitors, such as those profiled here, cannot activate NLRP1 in N-TERT cells. Their cytotoxic effects were likely caused by well-established pathways involving short-lived anti-apoptotic proteins such as MCL-1 (Figure 4E). Neither was the ability of UVB, ANS, DON and HYGRO to trigger NLRP1 caused by perturbed translation of the NLRP1 mRNA itself, since 1) none of these triggers have known sequence specificity, 2) both endogenous NLRP1 or overexpressed NLRP1 (with artificial UTRs and polyA signals) responded to ANS and UVB, and 3) UVB and ANS cannot activate NLRP1 orthologs and paralogs with similar domain arrangements (Figure 5A, S5A). ANS treatment under cytotoxic conditions (2μM, 5 hours) also had no effect on DPP9 enzymatic activity (Figure S4L). Altogether, these data provided further proof that ZAKɑ-activating ribotoxins, but not general translational inhibitors, specifically activate the NLRP1 inflammasome via a common pathway with UVB but distinct from those employed by VbP and dsRNA.

**Figure 5.**
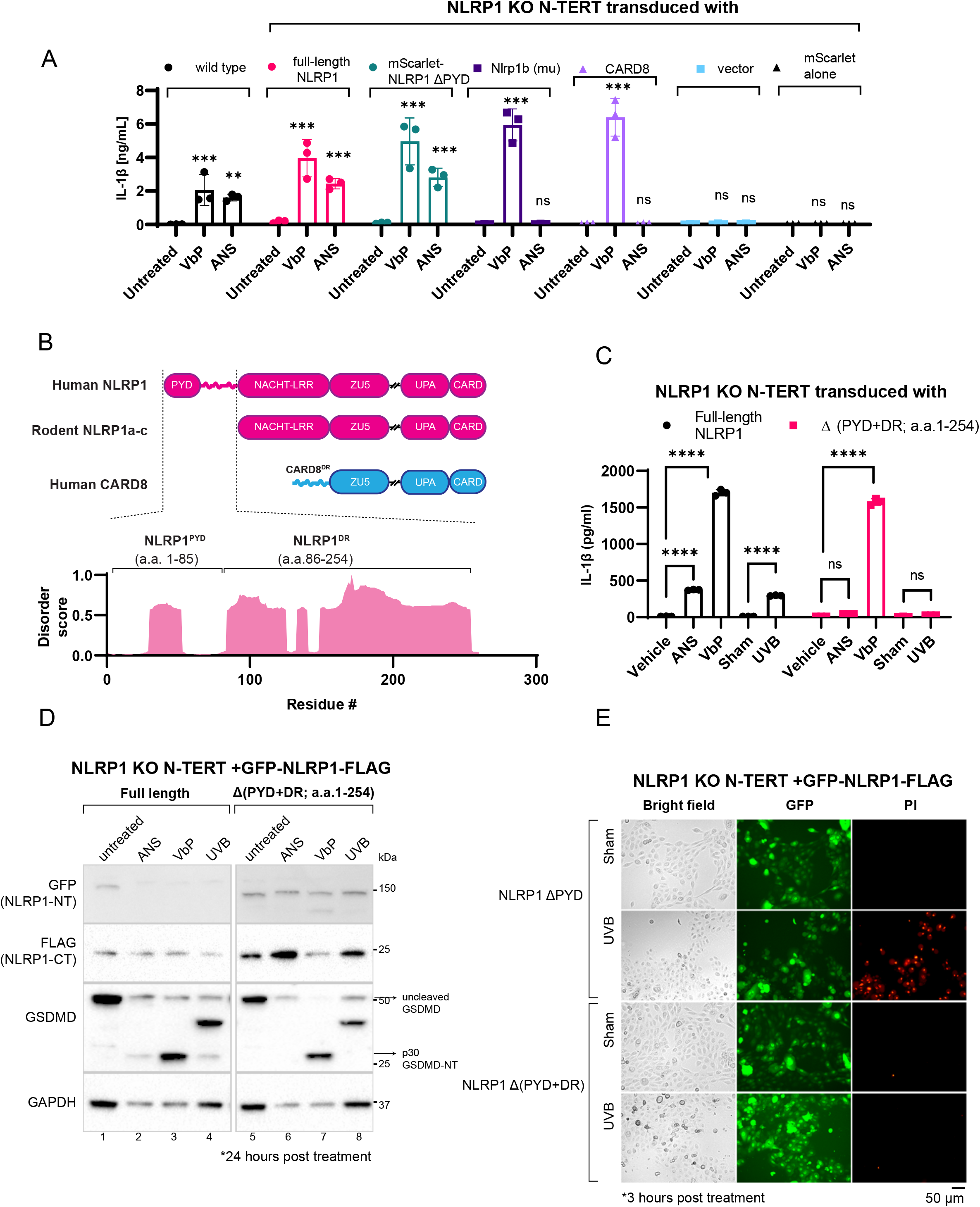
A human specific linker region is critical for ZAKɑ-dependent, but not VbP-dependent NLRP1 activation. A. IL-1β ELISA from *NLRP1* KO N-TERT cells reconstituted with the indicated NLRP1 variants and CARD8 treated with ANS. Note that this experiment was carried out independently in the Hornung laboratory in order to independently validate the results first obtained in the Zhong laboratory; hence a separate protocol involving a higher cell number was intentionally chosen to validate the differences seen between different sensors. B. Comparison between the domain structures of human NLRP1, rodent NLRP1a-c and human CARD8. The predicted disorder score was calculated for a.a. 1-300 of human NLRP1. C. IL-1β ELISA from *NLRP1* KO N-TERT cells reconstituted with GFP-full length NLRP1 or NLRP1 lacking PYD+DR (a.a.1-254). Cells were sham- or UVB-irradiated and harvested 24 hours post-treatment. D. Immunoblot from cells shown in C. E. PI and GFP fluorescence from UVB irradiated *NLRP1* KO N-TERT cells reconstituted with GFP-full length NLRP1 or NLRP1 lacking PYD+DR (a.a.1-254) grown in PI containing media. Images were acquired 3 hours after irradiation at 20x magnification.

Finally, to validate the regulation of NLRP1 by ZAKɑ in a more physiological setting, we tested direct chemical ZAKɑ activation on 3D organotypic skin. This experimental system was similar to those originally used to characterize UVB-triggered inflammasome activation in human skin (*22, 66*) (Figure 4H). In agreement with the N-TERT 2D culture experiments, both ANS- and VbP-treated organotypic skin displayed extensive epidermal dyskeratosis, i.e., abnormal keratinocytes with cytosolic vacuolization (marked by low eosin staining) and condensed, hematoxylin-rich nuclei (Figure 4J, red arrows). Immunostaining revealed GSDMD-NT p30 was prominently enriched at the cell membranes, especially around areas of dyskeratosis (Figure 4J, yellow arrows). These features were absent from untreated, as well as PURO-treated cultures, which displayed histologically different cell death features (Figure 4J, black arrows). Similarly, only ANS- and VbP-treated organotypic skins secreted significant amounts of IL-1β and IL-18 as compared to untreated cultures (Figure 4I). These results further proved that ZAK activation by ANS, similar to UVB, activates inflammasome-driven pyroptosis in a model of intact human skin.

Although skin epidermis likely represents the most biologically relevant tissue for the effect of UVB, ribotoxin exposure could cause systemic intoxication. For instance, DON, a fungal toxin, can cause acute toxic effects in people when ingested as part of contaminated cereal grains. Hence, we tested if ZAKɑ-activating toxins, exemplified by ANS, could activate NLRP1 in other human cell types. Using IL-1β secretion, GSDMD cleavage and ASC polymerization as surrogate markers for inflammasome activation, we found that ANS, similar to VbP, could induce inflammasome-driven pyroptosis in all NLRP1-expressing primary human cell types examined, including bronchial epithelial cells (NHBEs), foreskin keratinocytes and aortic endothelial cells (HAECs) (Figure S4N, S4O, S4P). HAECs stood out as particularly informative as it co-expressed NLRP1 and its ortholog CARD8 to high levels. We therefore knocked out *CARD8*, *NLRP1* and *CASP1* in HAECs. *NLRP1* and *CASP1* deletion abrogated GSDMD cleavage upon ANS treatment, whereas *CARD8* KO did not differ from control cells (Figure S4Q, lanes 3 vs 6, S4R). The lack of effect of ANS on CARD8 was further shown using MV-4-11 cells, which express CARD8 as the sole VbP-responsive inflammasome sensor. ANS did not lead to GSDMD cleavage in MV-4-11 cells (Figure S4S, S4T). These results demonstrate that ZAKɑ only controls the human NLRP1 but not the CARD8 inflammasome. Furthermore, the ZAKɑ dependent pathway of NLRP1 inflammasome activation operates in multiple primary human cell types, and is not restricted to N-TERT cells.

### The human-specific linker region (NLRP1^DR^) is necessary and sufficient to respond to ZAKɑ-activating agents

Next we wondered if UVB and ZAKɑ-activating agents could activate rodent NLRP1 molecules, which differ from human NLRP1 both in terms of protein structure and ligand specificity. Since murine epidermal keratinocytes express very low or undetectable levels of canonical inflammasome components, including Nlrp1b, Casp1 and Gsdmd (*23, 38*), mouse keratinocytes are not suitable to study the effect of UVB on the endogenous mouse Nlrp1b inflammasome. These observations also suggest that simple transgenic expression of human NLRP1 in murine skin cannot reconstitute a functional inflammasome response. In view of this, we reconstituted human NLRP1, human NLRP1 lacking the PYD domain (NLRP1 ΔPYD), murine NLRP1B (muNLRP1B) and human CARD8 in NLRP1 KO N-TERT cells, using a similar strategy previously used to study viral dsRNA-triggered NLRP1 activation (*20*) (Figure S5B). All overexpressed heterologous sensors restored VbP-induced IL-1β secretion, suggesting that they formed functional inflammasomes within N-TERT cells (Figure 5A). Furthermore, NLRP1 KO cells expressing muNLRP1B gained the ability to secrete IL-1β in response to anthrax lethal factor (LT), a rodent specific NLRP1B trigger that cannot activate human NLRP1 (Figure S5A). By contrast, ZAKɑ-activator ANS only induced IL-1β in cells rescued with human NLRP1. Thus, ANS, similar to dsRNA and enteroviral 3Cpro, is a specific trigger for human NLRP1 (Figure 5A). Additionally, like VbP and dsRNA, the non-signaling PYD of human NLRP1 is not involved in ANS response (Figure 5A).

Next we tested the effect of ANS on wild type- and mNLRP1a/b/c KO BMDMs. ANS triggered a small amount of IL-1β release, which was detectable by ELISA but not immunoblot, regardless of the NLRP1 genotype (Figure S5F, S5G). Therefore, the effect of ANS in murine BMDMs is not mediated by endogenous mNLRP1a/b/c. ANS has previously been reported to activate the NLRP3 inflammasome in murine BMDMs (*67*) (*68*). However, this effect was attributed to the global inhibition of translation, not to a single upstream regulator such as ZAKɑ. Therefore, these findings are not in conflict with the ZAKɑ-dependent activation of NLRP1 in human skin keratinocytes.

These results provided a strong clue to the molecular determinant that governs ZAKɑ-dependent NLRP1 activation. Human NLRP1 harbors a specific N-terminal extension encompassing the non-signaling PYD and an extended linker, which is absent in rodent NLRP1 orthologs and CARD8. The linker region is predicted to be intrinsically disordered (*69*), similar to the N-terminal linker of CARD8 (Figure 5B). Recent work by Chui and colleagues documented a critical role of CARD8 linker in VbP-dependent CARD8 inflammasome activation (*70*). Motivated by this discovery, we wondered if the NLRP1 linker region (a.a. 86-254, hereby termed NLRP1^DR^) also contributed to NLRP1 activation in response to either UVB, ANS or VbP. While removal of the PYD (a.a. 1 −86) did not impact NLRP1 activation, deletion of the NLRP1^DR^ selectively abrogated UVB- and ANS triggered NLRP1 activation, as evidenced from 1) the loss of IL-1β secretion (Figure 5C, S5E), 2) GSDMD cleavage (Figure 5D) and 3) PI uptake at early time points (Figure 5E). By contrast, VbP-dependent NLRP1 inflammasome activation was not affected. Similar results were obtained in 293T-ASC-GFP cells (Figure S5C, S5D). In addition, NLRP1^DR^ deletion diminished NT degradation (Figure 5D, Figure S5E, lane 1 vs lane 5; lane 2 vs lane 6), similar to the role of CARD8 linker region reported by Chui et al., (*70*). Although the deletion of NLRP1^DR^ also affected VbP-induced NT degradation, it did not affect VbP-triggered pyroptosis (Figure 5D). This was consistent with the recent finding that the disassembly of the NLRP1:DPP9 complex played a more dominant role in VbP-dependent human NLRP1 activation (*17, 18*). Taken together, our data demonstrate that NLRP1^DR^ is selectively required for ZAKɑ-dependent NLRP1 inflammasome activation.

Next we further dissected the function of NLRP1^DR^ by asking whether it would be sufficient to confer sensitivity to ZAKɑ-activating triggers in a heterologous context. To test this, we constructed a hybrid inflammasome sensor (termed NLRP1^DR^-CARD8^ZC^ with a C-terminal FLAG tag) by fusing NLRP1^DR^ to the signaling domains of CARD8 (ZU5-UPA-CARD) (Figure 6A, 6B lane 3). Since CARD8 itself is insensitive to UVB and ANS (Figure 6C, 6D lane 4 vs lane 9; S6B lane 4 vs lane 9), any neomorphic gain in inflammasome response to ANS and UVB could be attributed to NLRP1^DR^. When expressed in NLRP1 KO N-TERTs, full length CARD8 and NLRP1^DR^-CARD8^ZC^ both restored VbP-triggered IL-1β secretion (Figure 6E), GSDMD cleavage (Figure S6A lanes 4 and 5 vs lanes 9 and 10) and rapid PI uptake (Figure 6F), while only NLRP1^DR^-CARD8^ZC^-expressing cells displayed these hallmarks of inflammasome activation in response to ANS (Figure 6E, 6F, S6B, lane 10), UVB (Figure 6C, 6D, lane 10) and HYGRO (Figure S6C, S6D). These results provided further proof that NLRP1^DR^ is both necessary and sufficient for NLRP1 to be switched on by ZAKɑ-activating agents.

**Figure 6.**
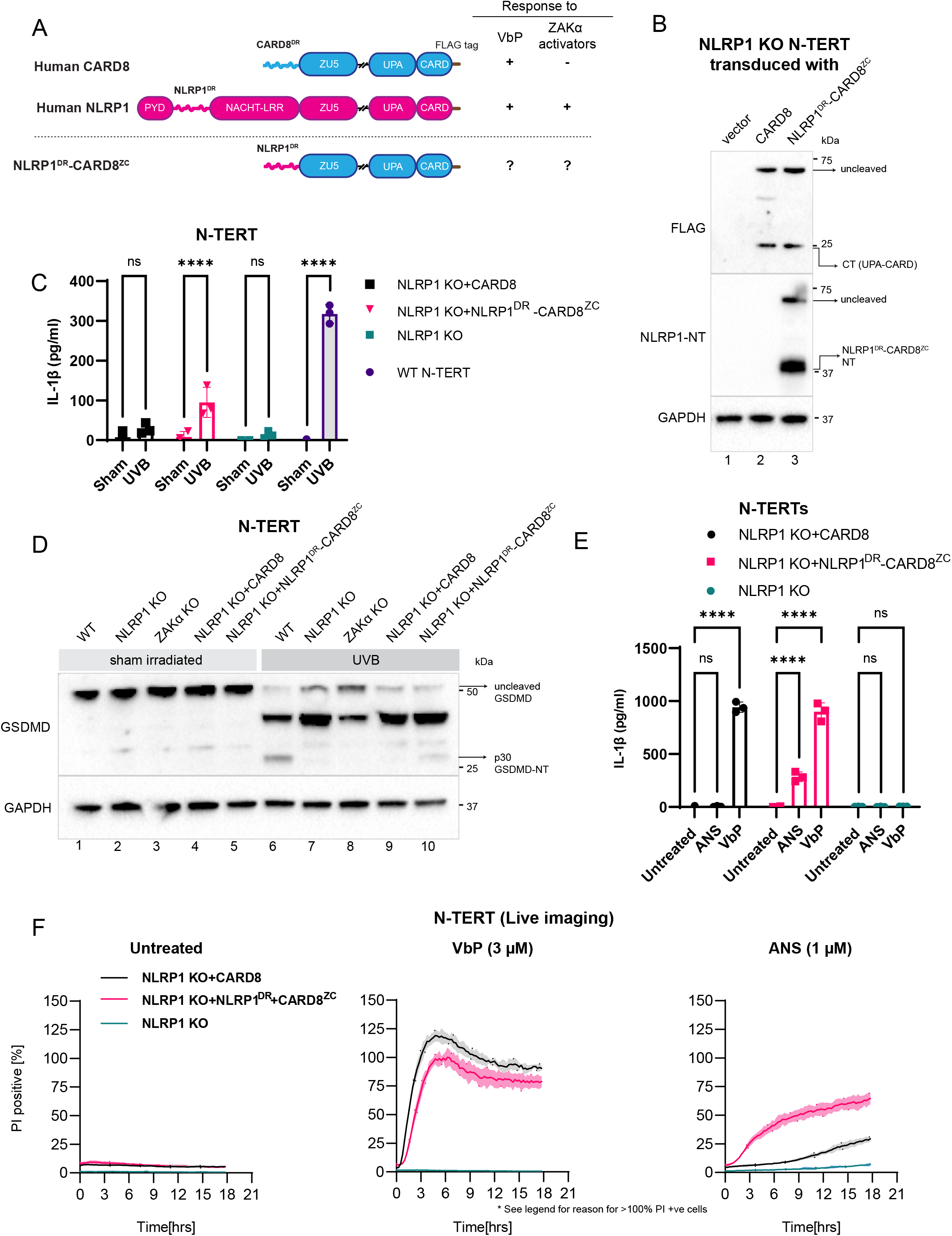
NLRP1^DR^ converts CARD8 to a UVB and ANS sensor. A. Comparison of the domain arrangements of human NLRP1 and CARD8, and the hybrid sensor NLRP1^DR^-CARD8^ZC^ B. Immunoblot of *NLRP1*KO N-TERT cells transduced with C-terminal FLAG tagged CARD8 and NLRP1^DR^-CARD8^ZC^. Note that the NLRP1 NT antibody recognizes NLRP1^DR^ and therefore detects NLRP1^DR^-CARD8^ZC^. C. IL-1β ELISA from sham- or UV-irradiated wild-type N-TERT cells or *NLRP1* KO cells transduced with CARD8 and NLRP1^DR^-CARD8^ZC^. Media were harvested 24 hours post irradiation. D. Immunoblot from the cells treated in C. The GSDMD Ab recognizes both full length and cleaved forms, including p43 and p30. E. IL-1β ELISA from N-TERT cells expressing the indicated sensors treated with 1 μM ANS or 3 μM VbP for 24 hours. F. Quantification of the percentage of PI positive *NLRP1* KO cells expressing CARD8 or NLRP1^DR^-CARD8^ZC^ during 18-hour incubation of ANS or VbP. For CARD8 expressing cells, nearly all cells underwent pyroptosis at around 4-5 hours. Thereafter the disintegrated pyroptotic corpses detached en masse from plates, causing an artifactual increase in PI positive cells above 100%. This issue did not affect the comparison of molecular readouts of pyroptosis at early time points (∼3 hours) used as an inflammasome readout for ZAKɑ throughout this study.

### ZAKɑ-activating agents cause NLRP1^DR^ hyperphosphorylation

We next tested the most parsimonious hypothesis: NLRP1^DR^ harbors a ZAKɑ-dependent degron that destined it for proteasomal degradation. However, this did not seem to be the case, as NLRP1^DR^ as a GFP fusion protein (NLRP1^DR^-GFP) in fact showed increased fluorescence 24 hours after ANS treatment (Figure S7A). Thus, although NLRP1^DR^ contributes to NLRP1 NT degradation, other structural elements on NLRP1 and/or cellular factors are necessary. Nonetheless, in the course of this analysis, we noticed a marked bandshift for NLRP1^DR^-GFP on immunoblot whenever the cells were treated with UVB or ANS. This bandshift was sensitive to post-lysis treatment with lambda phosphatase (LPP), confirming that it was caused by phosphorylated NLRP1^DR^ (Figure S7B). As a negative control, NLRP1^PYD^-GFP did not demonstrate any bandshift, arguing against the possibility that ANS and UVB treatment indiscriminately phosphorylate all overexpressed proteins. Next we examined the phosphorylation pattern of NLRP1^DR^ in greater resolution using PhosTag-containing SDS-PAGE. This experiment revealed that NLRP1^DR^ was in fact significantly phosphorylated in unstimulated cells, and became further phosphorylated by ANS and UVB (Figure 7A). Almost no unphosphorylated species was left 2 hours post treatment. We hereby refer to NLRP1^DR^ phosphorylation in unstimulated cells as ‘*basal phosphorylation*’ and ANS- or UVB-dependent NLRP1^DR^ phosphorylation as ‘*hyperphosphorylation’*. Importantly, other ZAKɑ-activating drugs, such as HYGRO and DON, also led to NLRP1^DR^ hyperphosphorylation, but not VbP or any of the non-ZAKɑ-activating cytotoxic compounds (Figure 7B). In *ZAKɑ* KO N-TERT cells, ANS- and UVB-induced NLRP1^DR^ ‘hyperphosphorylation’ was completely abrogated, while NLRP1^DR^ basal phosphorylation remained unaffected (Figure 7A, lane 3 vs. lane 4; lane 7 vs. 8). Thus ZAKɑ-dependent NLRP1^DR^ hyperphosphorylation occurs whenever NLRP1 is activated by UVB or ZAKɑ-activating toxins.

**Figure 7.**
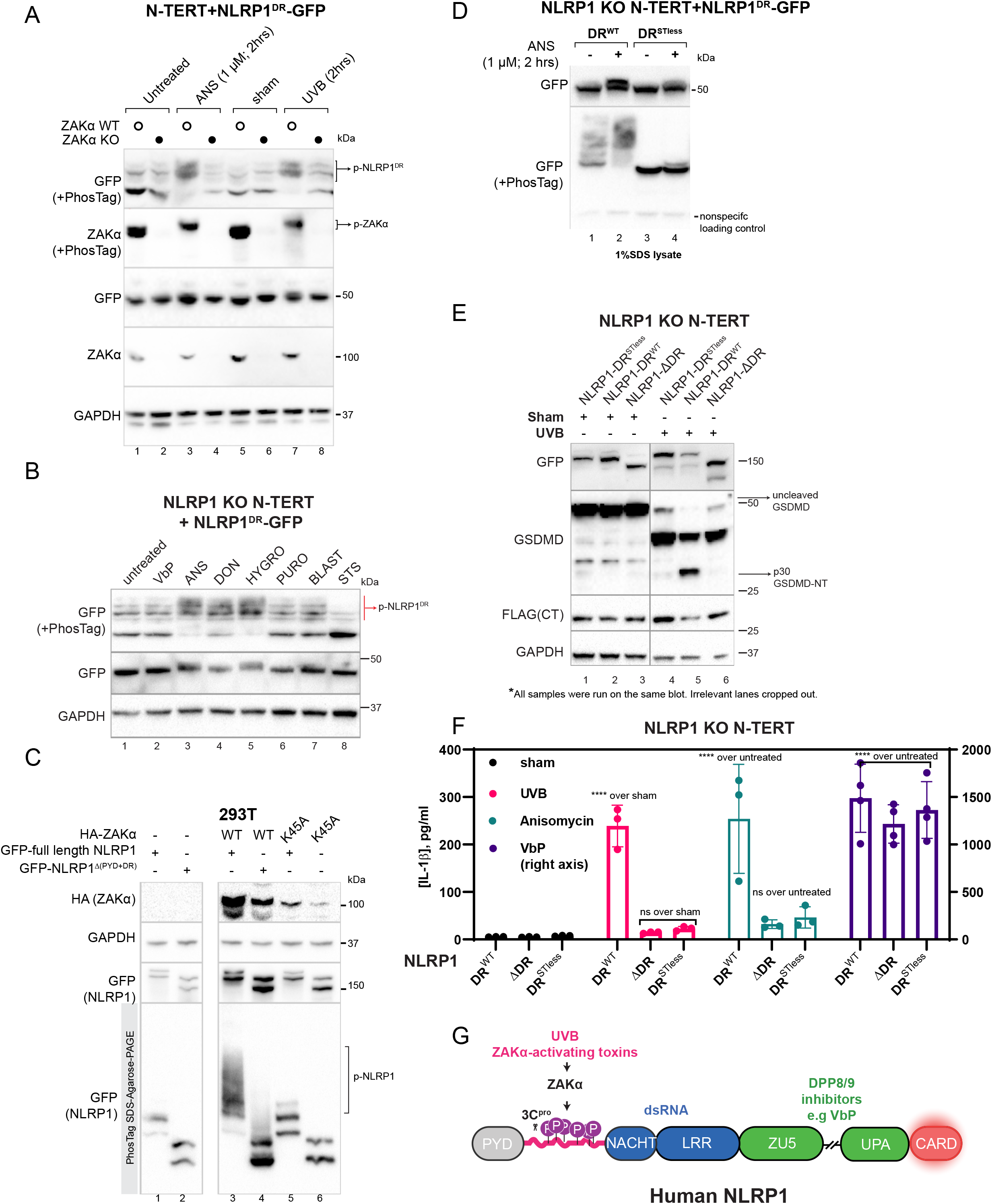
ZAKɑ-dependent NLRP1^DR^ hyperphosphorylation is required for UVB and ribotoxin sensing by human NLRP1. A. Immunoblot following SDS-PAGE or PhosTag SDS-PAGE of wild-type and *ZAKɑ* KO N-TERT cells expressing NLRP1^DR^-GFP. Cells were harvested 2 hours post ANS treatment or UVB irradiation. B. GFP immunoblot of NLRP1^DR^-GFP in *NLRP1* KO N-TERT cells treated with the indicated drugs. Cells were harvested 3 hours post treatment. C. Immunoblot of GFP-tagged NLRP1 or HA-tagged ZAKɑ expressed in 293T cells following SDS-PAGE or PhosTag-SDS-Agarose-PAGE. Cells were harvested 48 hours post transfection. Note that it is common for phospho-species to appear as smears on PhosTag-SDS-Agarose-PAGE (see also Figure S7C). D. GFP immunoblot of DR mutants expressed in *NLRP1* KO N-TERT cells following SDS-PAGE and PhosTag SDS-PAGE E. GFP, FLAG and GSDMD immunoblot of *NLRP1* KO N-TERT cells expressing the indicated NLRP1^ΔPYD^ DR mutants. All constructs were fused with GFP at the N-terminus and FLAG at the C-terminus; therefore GFP reports on NLRP1-NT, while FLAG indicates NLRP1-CT. GSDMD p30 is marked as GSDMD-NT with black arrow. Cells were harvested 24 hours post UVB treatment. F. IL-1β ELISA from *NLRP1* KO N-TERT cells expressing the indicated NLRP1^ΔPYD^ DR mutants 24 hours after UVB, ANS or VbP treatment. G. Graphical representation of the ‘division of labor’ between NLRP1 domains with regard to known human NLRP1 agonists.

Next we sought orthogonal proof that activated ZAKɑ causes hyperphosphorylation of full length NLRP1 in the DR region. We were unable to reliably detect endogenous NLRP1-NT in N-TERT cells using PhosTag SDS-PAGE, as all commercial NLRP1-NT antibodies (R&D Systems, #AF6788 and Biolegend #9F9B12) recognize epitopes within NLRP1^DR^ (potential gain or loss of hyperphosphorylation might mask the epitopes). As an alternative, we visualized overexpressed full-length NLRP1 in 293T cell lysates using PhosTag-containing SDS-agarose-PAGE, a technique designed for the detection of phosphorylation on large proteins (>150 kDa). While NLRP1 by itself migrated as two discrete bands (corresponding to uncleaved NLRP1 and NLRP1-NT), co-expression of wild-type, kinase-active ZAKɑ induced a large bandshift, which is indicative of extensive phosphorylation of full length NLRP1 (Figure 7C, lane 3). Importantly, deletion of the DR region largely abrogated NLRP1 phosphorylation (Figure 7C, lane 4), as did a catalytic mutation (p.K45A) in the ZAKɑ kinase domain and the removal of ZAKɑ RSR sensing domains (CTD and S domains) (Figure 7C, lanes 5-6, Figure S7C, lanes 2 vs 4). Taken together, these results demonstrate that ZAKɑ-activating agents, such as ANS and UVB induce phosphorylation of full length NLRP1 primarily within the disordered linker region.

The remarkable extent of ZAKɑ-dependent NLRP1^DR^ phosphorylation is consistent with the high number of serines and threonines in this region (25 serines and 15 threonines among 169 amino acids). In particular, serine and proline residues, which are commonly found in kinase recognition motifs at the P1-P2 positions, are both over-represented in NLRP1^DR^ (Figure S7D) To test the functional consequence of hyperphosphorylation in NLRP1 inflammasome signaling, we mutated all the serines and threonines within NLRP1^DR^ to alanines (referred to as ‘STless’). This mutant largely abrogated NLRP1^DR^ basal phosphorylation and hyperphosphorylation after ANS and UVB treatment (Figure 7D). When expressed in *NLRP1* KO N-TERT cells, NLRP1-STless restored the VbP-triggered pyroptosis to wild-type levels, but remained completely unresponsive to UVB and ANS by all measures tested, including IL-1β secretion (Figure 7F), GSDMD p30 cleavage (Figure 7E, lane 4 vs lane 5), and rapid PI uptake (Figure S7E) as well as NLRP1-NT degradation (Figure 7E, lane 4 vs lane 5, top panel). Based on these results, we conclude that ZAKɑ-dependent NLRP1^DR^ hyperphosphorylation is required for UVB- and ANS-triggered NLRP1 activation, but is not required for activation of NLRP1 by DPP8/9 inhibition.

## DISCUSSION

For many years it was known that UVB irradiation, the most common cause for sunburn, leads to caspase-1-dependent pyroptotic cell death and IL-1β secretion in human skin keratinocytes (*22, 66, 71*). Initially thought to be an NLRP3-associated phenomenon, the Beer group and colleagues provided concrete proof using CRISPR/Cas9-knockout cells that NLRP1, instead of NLRP3, is the primary inflammasome sensor for UVB. However, the mechanisms of UVB sensing by NLRP1 remained unknown. In this work, we closed the knowledge gap by resolving key events in UVB-triggered NLRP1 inflammasome activation in keratinocytes. By inducing cellular RNA photo-lesions that stall the ribosomes, UVB activates the ribotoxic response (RSR) kinase ZAKɑ in a cell-autonomous manner. Auto-phosphorylated, activated ZAKɑ then acts as phosphorylation-dependent ‘ON’ switch for NLRP1 that operates independently of DPP8/9. In addition, we showed that known and novel ZAKɑ-activating microbial toxins all function as specific activators for human, but not rodent NLRP1. Unexpectedly, the cause for this species specificity was traced to hyperphosphorylation events on a previously uncharacterized, disordered linker region that is only found in human NLRP1. Transplanting this linker region to the signaling domains of CARD8 created a minimal UVB inflammasome sensor. Based on these findings, we propose a new pathway of NLRP1 activation controlled by ZAKɑ and NLRP1^DR^ hyperphosphorylation. Our discoveries markedly expanded the repertoire of known inflammasome activators for primary human cells, including not only skin keratinocytes, but also airway epithelial cells and endothelial cells. We also uncovered a ‘division of labor’ amongst the discrete NLRP1 domains, with each domain involved in recognizing a distinct trigger (Figure 7G).

### NLRP1 inflammasome as a new RSR effector: implications for anti-microbial immunity

Although a detailed dissection of the RSR pathway itself falls outside of the scope of this study, the seminal work by Wu et al., Vind et al., and others provided strong proof that UVB irradiation and a subset of ribotoxins activate ZAKɑ by stalling elongating ribosomes (*56, 57*). It is interesting to speculate that ZAKɑ, via its ribosome binding domains, can recognize a particular structural conformation associated with a single stalled ribosome, or two, or a series of collided ribosomes. Our work adds to the growing body of literature that demonstrates that ribosomes themselves serve as signaling hubs that can give rise to extremely specific cellular outcomes (e.g. RQC, ISR vs RSR) (*58*) (*54, 59*), each defined by a dedicated set of sensors and effectors. In this regard, care must be taken not to equate pleiotropy with nonspecificity. No doubt, ZAKɑ-activating agents such as UVB and the ribotoxins characterized in this study have multiple cellular effects, but this does not preclude some of them from being highly specific. Throughout the course of this study, we used multiple specific inflammasome readouts (e.g. IL-1β p17 secretion, GSDMD p30 cleavage, PI uptake at early time points) coupled with genetic knockouts to demonstrate the specificity of ZAKɑ-dependent pathway of NLRP1 inflammasome activation. Wherever possible, we also distinguished the effects of UVB and other ZAKɑ-activating ribotoxins on NLRP1 from their non-inflammasome related effects. Importantly, our analyses established that the ability of UVB and certain ribotoxins to activate human NLRP1 is not related to global translational inhibition or apoptotic cell death. Therefore, we conclude the ZAKɑ-activating agents, as exemplified by UVB, ANS, HYGRO and DON, represent a new class of bona fide NLRP1 inflammasome activators in human cells.

Another key implication of our findings is that NLRP1 can be thought of as a new effector of the RSR pathway. Conversely, ZAKɑ-dependent RSR qualifies as innate immune sensing pathway that responds to toxin- or irradiation-triggered perturbation of an essential host process, i.e., elongation of translating ribosomes. This pathway likely functions in parallel with others that can respond to translational inhibition, such as those involving anti-apoptotic protein MCL-1 triggered by viral proteins (*72*). Why do human cells evolve multiple innate immune pathways to sense perturbation of translation? Further investigation is no doubt required, but one tantalizing clue lies in the fact that nearly all known ZAKɑ-activating ribotoxins target the *elongation* step of translation and are of bacterial and fungal origin, whereas nearly all viral inhibitors of translation target the *initiation* step of translation. Thus, it is tempting to speculate the ZAKɑ-NLRP1 pathway specifically defends against bacteria/fungi whereas the MCL-1-dependent pathway specializes in viral immunity (*72*). Germline ZAKɑ mutations are also implicated in Mendelian disorders in humans; it will be interesting to examine if aberrant RSR signaling and/or inflammasome regulation contribute to the pathogenesis of these conditions (*64, 73*).

### Species-specific role of the NLRP1 inflammasome in human skin

By delineating the molecular events of UVB-induced keratinocyte pyroptosis, our results also potentially explains the unique role of NLRP1 in human skin that is not shared by other inflammasome sensors. Germline NLRP1 mutations cause a spectrum of Mendelian disorders characterized by epithelial inflammation and hyperplasia (*34–36*). Certain NLRP1 single nucleotide polymorphisms (SNPs), some of which are found in NLRP1^DR^, increase risk for auto-immune diseases such as psoriasis and vitiligo (*37, 74*). It is likely that these SNPs lead to an altered inflammasome response in response to sunburn or ribotoxin exposure. UVB-associated skin damage is also well known to precipitate flares of auto-immune diseases such as systemic lupus erythematosus and bullous pemphigoid (*75*). Our findings should give fresh impetus to the development of pharmacologic NLRP1 or ZAKɑ inhibitors for the treatment of these conditions.

### What is the direct kinase that hyper-phosphorylates NLRP1^DR^?

Our data could not thus far identify the direct kinase that hyper-phosphorylates NLRP1^DR^, be it ZAKɑ itself or another kinase downstream of it. It is also likely that multiple RSR-dependent kinases act in concert, each phosphorylating a subset of serine and threonine residues spread across the entire NLRP1^DR^. We also inadvertently discovered that NLRP1 is phosphorylated at multiple sites in unstimulated N-TERT cells by non-ZAKɑ kinases. The functional consequences of this await further investigation. Our attempts at mapping the exact phosphorylation sites on NLRP1^DR^ were unsuccessful due to low coverage of the region. As Vind et al., and Wu et al., established that the kinase activity of ZAKɑ must be unleashed by stalled or collided ribosomes, NLRP1^DR^ phosphorylation is unlikely to be reconstituted in a simple *in vitro* kinase assay of recombinant ZAKɑ and NLRP1. This was also supported by our observation that neither ZAKβ, which shares an identical kinase domain as ZAKɑ, nor ZAKɑ ΔCTD, which retains the full kinase domain but cannot bind to the ribosomes, could fully phosphorylate NLRP1. Hence additional cellular factors, putatively a fully assembled but stalled ribosome complex, must be required. At the time of writing, *in vitro* reconstitution of the ZAKɑ activation reaction by purified stalled/collided ribosomes have not been achieved.

### How does NLRP1^DR^ control N-terminal fragment degradation?

Similar to VbP, dsRNA and enteroviral proteases, ZAKɑ-activating agents induce NLRP1 NT degradation. When the linker region is disrupted, the resulting ΔDR or NLRP1-DR^STless^ mutant no longer underwent NT degradation and could no longer sense UVB and other ZAKɑ-activating agents. Although we initially hypothesized that NLRP1^DR^ harbored a direct phospho-degron, it did not undergo proteasomal degradation by itself, at least when expressed as a GFP-fusion protein. Hence, additional cellular factors or structural elements on NLRP1 must be required downstream to trigger NLRP1 activation. Alternatively, the substantial shift in charge caused by NLRP1^DR^ hyperphosphorylation might cause a conformational change in NLRP1 NT directly. With the exception of enteroviral 3C proteases (*19*), the exact cellular factors governing human NLRP1 NT (or CARD8 NT) ‘functional degradation’ have not been identified, even for the most well established NLRP1/CARD8 agonist, VbP. Although a detailed characterization of NLRP1 NT degradation falls outside of the scope of the current study, it is tempting to speculate that each NLRP1 trigger employs a different route of NLRP1 NT degradation. Our findings here should provide a framework for future work investigating this hypothesis.

## ACKNOWLEDGMENTS

The authors would like to thank the SRIS Asian Skin Biobank (ASB), especially Aishah Alimat, Joycelyn Lee and Alicia Yap, for generating and providing human primary skin cells. ASB is funded by Singapore’s Agency for Science, Technology & Research (A*STAR) through core fund and under the IAF-PP Project (H1701a0004). The authors are especially indebted to Prof. Bruno Reversade, Dr. Kenneth Lay and Dr. Shifeng Xue for their detailed critique of this manuscript.

## AUTHOR CONTRIBUTIONS

F.L.Z. conceived of and supervised this study. K.S.R. designed and performed experiments, analyzed the data and co-wrote the manuscript, with significant contribution from G.A.T., P.R., S.B., Z.S., S.B.. S.B. performed the N-TERT constitution experiments and analyzed the data with supervision from V. H. C.R.H. performed the BMDM experiments with supervision from S.L.M. K.W.C. performed independent validation experiments with BMDM. R.N. and W.C. performed all endothelial cell experiments with supervision from L. H. C.K.L. performed cytokine analysis with supervision from J.E.C. C.B. and the Asian Skin Bank assisted with primary cell derivation and culture. K.Y.T. carried out all mass spec. experiments with supervision from R.S. All co-authors contributed to writing the manuscript.

## METHODS

### Cell culture and chemicals

293Ts (ATCC #CRL-3216), MV-4-11 (ATCC #CRL-9591), mBMDMs (were a kind gift from Linfa Wang, Duke-NUS, Singapore) and normal bronchial epithelial cells (NHBE, Lonza #CC-2541) were cultured according to manufacturer’s protocols. Immortalised human keratinocytes (N/TERT-1 or N-TERT herein) were provided by H. Rheinwald (MTA) (*76*). Primary human keratinocytes were derived from the skin of healthy donors and obtained with informed consent from the Asian Skin Biobank (ASB) (https://www.a-star.edu.sg/sris/technology-platforms/asian-skin-biobank). Primary endothelial cells (HAoEC-c or HAEC herein), excised from the ascending and descending aortic arch were purchased from Promocell (#C-12271). All cell lines underwent routine mycoplasma testing with MycoGuard (Genecopoeia #LT07-118). The following drugs and chemicals were used as part of this study: Staurosporine (STS, MCE, #HY-151141), Puromycin (PURO, Sigma, #P9620), Talabostat (VbP, MCE, #HY-13233), Anisomycin (ANS, MCE, #HY18982), Etoposide (EPEG, MCE, #HY13629), Camptothecin (CPT, MCE, #HY16560), Blasticidin (BLA, Sigma, SBR00022), Geneticin (G418, Sigma #G8168), Hygromycin B (HYGRO, Thermo Scientific, 10687010), Belnacasan (MCE, HY-13205), MCC950 (MCE, #HY-12815A), Cisplatin (MCE, HY-17394), Nilotinib (MCE, HY-10159A), Nigericin (Sigma, N7143), Emetine (EME, MCE, HY-B1479B), Harringtonine (HTN, MCE, HY-N0862), 5-BrDU (BrDU, MCE, HY-15910), 4-Thiouridine (4SU, Sigma, T4509), Poly I:C (Invivogen, tlr1-pic). Bortezomib (BTZ) and MLN9424 provided by D. Lane, p53 lab, ASTAR, Singapore. For mouse studies, 1 mg/ml anthrax lethal factor (List Biological Labs #LL-169B) and 1 mg/ml protective antigen (List Biological Labs #LL-171E) for anthrax lethal factor (LF) was used.

### UVB and UVA irradiation

For UVA and UVB irradiation experiments, cells were seeded on 6 cm dishes, incubated for 24 hours and washed once in phosphate buffered saline pH 7.4 before being exposed to indicated dose of irradiation using a BIO SUN microprocessor controlled, cooled UV irradiation system (BIO-SUN, Vilber). After exposure, PBS was replaced by keratinocyte medium and cells were incubated for indicated time.

### Lambda Phosphatase dephosphorylation assay

293T cells were transfected with either the NLRP1 PYD fragment(a.a. 1-85) or NLRP1 DR fragment(a.a. 86-254) tagged with GFP. After 2 days, cells were treated with ANS for 3 hours, and subsequently harvested and lysed in tris-buffered saline 1% NP-40 with protease inhibitors (Thermo Scientific, #78430). Protein concentration was determined using the Bradford assay (Thermo Scientific, #23200). For each reaction, 40 μg of protein lysate topped up to 40μl with distilled water was added with 5μl of 10X NEBuffer for Protein MetalloPhosphatases (PMP) and 5μl of 10 mM MnCl_2_ to make a total reaction volume of 50μl. 1μl of Lambda Protein Phosphatase (NEB, P0753S) was added and samples were incubated at 30°C for 30 minutes. Final reaction products were prepared for SDS-PAGE immunoblotting using the protocol mentioned below.

### Cytokine analysis

To measure secreted cytokine and chemokine levels a human IL-1β enzyme linked immunosorbent assay (ELISA) kit (BD, #557953), human IL-6 ELISA kit (R&D Systems, D6050) and a human IL-8/CXCL8 ELISA kit (R&D Systems, D8000C) was used.

### Plasmids and preparation of lentiviral stocks

293T-ASC-GFP, N-TERT-ASC-GFP, N-TERT NLRP1 KO, N-TERT CASP1 KO, N-TERT ASC KO cells were previously described (*14*). All expression plasmids for transient expression were cloned into the pCS2+ vector backbone and cloned using InFusion HD (Clonetech). Constitutive lentiviral expression was performed using pCDH vector constructs (System Biosciences) and packaged using third generation packaging plasmids. ZAKɑ plasmids for expression in 293Ts were gifts from Simon Bekker-Jensen (Addgene plasmids # 141193, # 141194, # 141195, # 141196, #141197; http://n2t.net/addgene:141193; RRID:Addgene_141193).

### CRISPR-Cas9 knockout

Lentiviral Cas9 and guide RNA plasmid (LentiCRISPR-V2, Addgene plasmid #52961) was used to create stable deletions in N-TERT keratinocytes. The sgRNAs target sequences (5’ to 3’) are: NLRP1 (GATAGCCCGAGTGCATCGG), or NLRP1 (5′-GGAGTTAAGAGGGTGTCTGG-3′′), ASC (GCTAACGTGCTGCGCGACAT), CASP1 (ACAGACAAGGGTGCTGAACA), NLRP3 (GAATCCCACTGTGATATGCC), GSDMD (AGGTTGACACACTTATAACG), MAP3K20(ZAKɑ) sg1 (TGTATGGTTATGGAACCGAG), MAP3K20(ZAKɑ) sg4 (TGCATGGACCGGAAGACGATG), DDB2 sg2 (TGTAGCCCTCCTGTCAAAGG), DDB2 sg3 (CCCAACTCACCCCAGCACCG), ZNF598 (CGGCACTCGCGCCGGAACGA). Knockout efficiency was tested by immunoblot. Alternatively, Sanger sequencing of genomic DNA and overall editing efficiency were determined using the Synthego ICE tool (Synthego Performance Analysis, ICE Analysis. 2019. v2.0. Synthego, https://ice.synthego.com/#/).

### Immunoblotting

The following antibodies were used in this study: cleaved PARP1 (Abclonal, #WH162766), Cleaved CASP3 (Cell Signaling Technology, #9664), Full length GSDMD-FL (Abcam, #ab210070, ab209845), IL1β p17 specific (Cell Signaling Technology, #83186S), pro–IL-1β (AF-401-NA, R&D), CARD8 (Abcam, CARD8-NT: ab19485, CARD8-CT: ab241186), HA tag (Santa Cruz Biotechnology, #sc-805), GFP (Abcam, ab290), GAPDH (Santa Cruz Biotechnology, #sc-47724), ASC (Adipogen, #AL-177), CASP1 (Santa Cruz Biotechnology, #sc-622), caspase-1 p20 (1:1000; casper-1, AdipoGen) IL1β (R&D systems, #AF-201), FLAG (SigmaAldrich, #F3165), NLRP1 (R&D systems, #AF6788), NLRP1 (Biolegend #9F9B12),, cleaved GSDMD-NT (Asp275) (Cell Signaling Technology, #36425), GSDMDC1 (Novus Biologicals, NBP2-33422), DDB2 (Abcam, ab181136), phospho-p38 (Cell Signaling Technology, 4511T), phospho-JNK (Cell Signaling Technology, 4668T), ZAK (Bethyl Laboratories, A301-993A), MCL-1 (Abcam, ab32087), anti-thymine dimer antibody (Sigma, T1192). All horseradish peroxidase (HRP)– conjugated secondary antibodies were purchased from Jackson Immunoresearch (goat anti-mouse IgG: 115-035-166; goat anti-rabbit IgG: 111-035-144; and donkey anti-sheep IgG: 713-035-147). Blue-Native PAGE was carried out using th Native-PAGE system (ThermoFisher) with 10-20μg of total lysate. For SDS-PAGE using whole cell lysates, cells were resuspended in tris-buffered saline 1% NP-40 with protease inhibitors (Thermo Scientific, #78430). Protein concentration was determined using the Bradford assay (Thermo Scientific, #23200) and 20 μg of protein loaded, apart from cleaved GSDMD-NT visualisation where 40μg of protein was used. All primary antibodies were used at 250ng/ml. Visualisation of ASC oligomerization was previously described (*19*). For analysis of IL-1β in the media by immunoblotting, samples were concentrated using filtered centrifugation (Merck, Amicon Ultra, #UFC500396). Protein samples were run using immunoblotting, and then visualized using a ChemiDoc Imaging system (Bio-Rad). PhosTag SDS-PAGE was carried out using homemade 10% SDS-PAGE gel, with addition of Phos-tag Acrylamide (Wako Chemicals, AAL-107) to a final concentration of 30uM and manganese chloride(II) (Sigma-Aldrich, #63535) to 60uM. PhosTag-SDS-Agarose-PAGE gels were made to 3% polyacrylamide and 0.5% agarose with a final concentration of Phos-tag Acrylamide and MnCl_2_ as mentioned above.. Cells were directly harvested using Laemmli buffer, lysed with an ultrasonicator, and loaded into the Phos-tag gel to run. Once the run was completed, the polyacrylamide gel was washed in transfer buffer with 10mM EDTA twice, subsequently washed without EDTA twice, blotted onto 0.45um PVDF membranes (Bio-rad), blocked with 3% milk, and incubated with primary and corresponding secondary antibodies.

### Propidium iodide inclusion assay

N-T TERT cells of various genotypes were seeded at a cell density of 10000 cells/well in a black 96 well plate (PerkinElmer, CellCarrier-96 Ultra, #6055300). The next day cells were treated with chemicals and stained with 1μg/ml dilution of propidium iodide (PI, Abcam #ab14083) before observing the cells on a high content screening microscope (Perkin Elmer Operetta CLS imaging system) over 18 hours, capturing brightfield and fluorescent images every 15 minutes. Images were then stored and analyzed using the Harmony software (Version 6). For 5 fields of view per well, with 3 wells per treatment, the ratio of PI positive cells over normal cells was calculated. The number of live cells per field was counted using digital phase contrast images which can identify cell borders, while the number of PI stained regions identified through the PI channel (536nm/617nm) was counted as PI positive cells.

### Microscopy

Images of ASC-GFP specks were acquired in 3 random fields in 4’,6-diaminidino-2-phenylindole (DAPI, 358nm/461) and GFP (469nm/525nm) channels using the EVOS microscope (FL Auto M5000, #AMF5000) according to the manufacturer’s protocol. Quantification method of ASC-GFP specks was previously described in detail (*19*)

### DPP8/9 activity assay

293T cells were transfected with vector or wild-type DPP9. DPP9-transfected cells were treated with VbP, ANS or HTN. Cells were lysed in PBS 1% Tween-20, 48h after transfection and treatments. 0.3 μg of total lysate was then incubated with 0.1 μM uGly-Pro-AMC fluorescence substrate. AMC fluorescence was measured after 30mins at 25°C in a 50 μl reaction every minute on a spectrometer and the rate of Gly-Pro-AMC hydrolysis per minute calculated.

### Statistical analysis

Statistical analyses were performed using Prism 8 or 9 (GraphPad). The methods for statistical analysis were included in the figure legends. Error bars show mean values with SEM.

## Figures

**All significance values were calculated based on one-way ANOVA from three biological replicates, with each treatment/transfection considered a single replicate. Significance values were indicated as: *n.s* (non significant), \*\*\**P<*0.01, \*\*\**P<*0.001, \*\*\*\**P<*0.0001**

---

**Figure S1.**
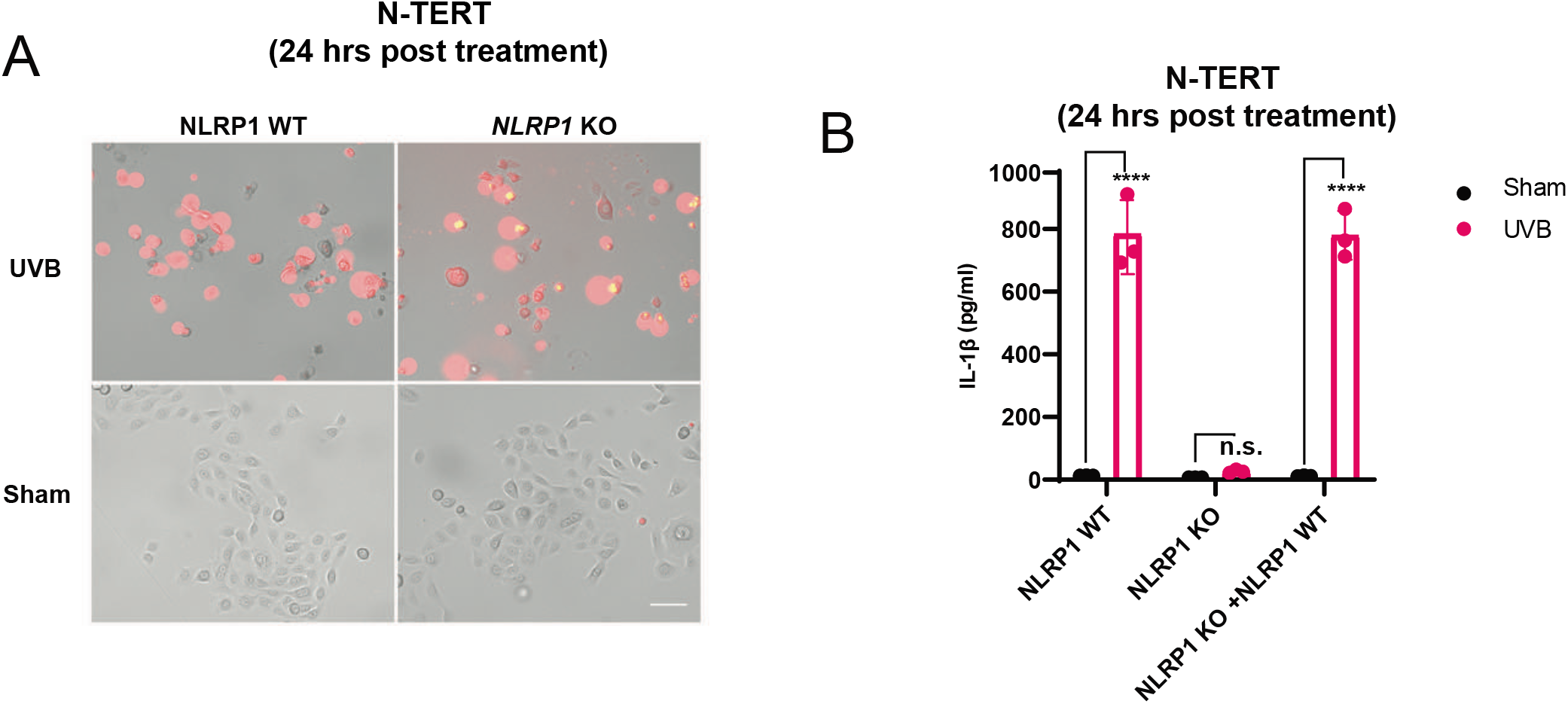
Further supporting evidence that UVB specifically activates human NLRP1. A. Representative images of PI positive wild type N-TERT cells or *NLRP1* KO N-TERT cells 24 hours after UVB (100mJ/cm^2^) or sham irradiation. B. IL-1β ELISA of N-TERT cells of the indicated genotypes. Cells were either sham or UVB (100mJ/cm^2^) irradiated and cell media collected 24 hours later.

**Figure S2.**
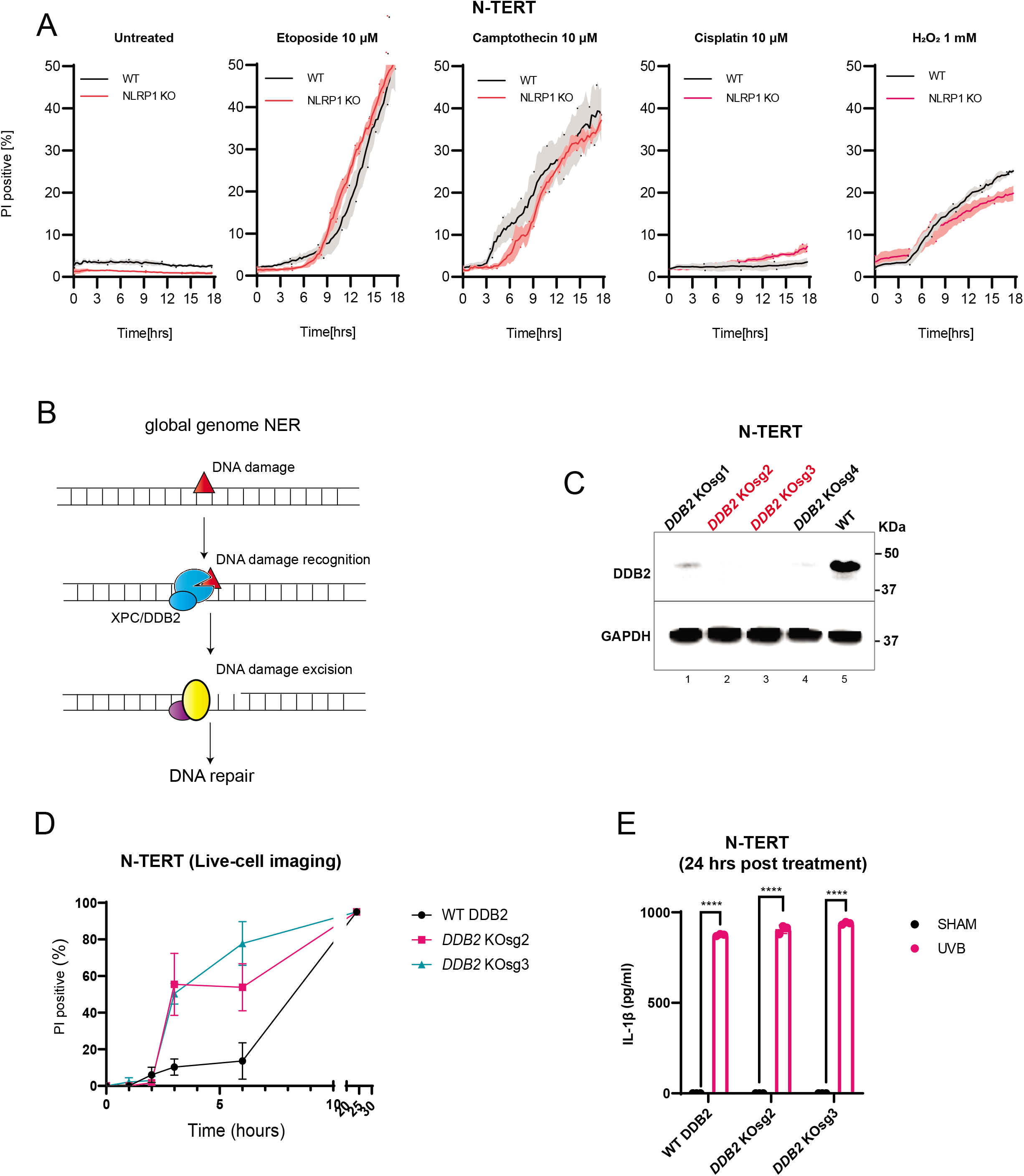
Additional evidence to support DNA independent response of UVB. A. Quantification of the percentage of PI positive N-TERT cells during an 18-hour incubation with the indicated drugs. Cells were grown in 0.25 μg/mL PI and imaged at 15 min intervals. B. Outline of the NER pathway. C. Immunoblot validation of WT or *DDB2* KO N-TERT using guide RNA 1-4. The *DDB2* KO used for further study is shown in red. D. Percentage of PI positive wild type N-TERT cells or *DDB2sg2/DDB2sg3* KO N-TERT cells following UVB (100mJ/cm^2^) irradiation. Cells were imaged after 2,3,4,5,6 and 24 hours after UVB irradiation. E. IL-1β ELISA of WT or *DDB2 sg2/DDB2 sg3* KO N-TERT after sham or UVB (100mJ/cm^2^) irradiation. Cell culture media were collected 24 hours later.

**Figure S3.**
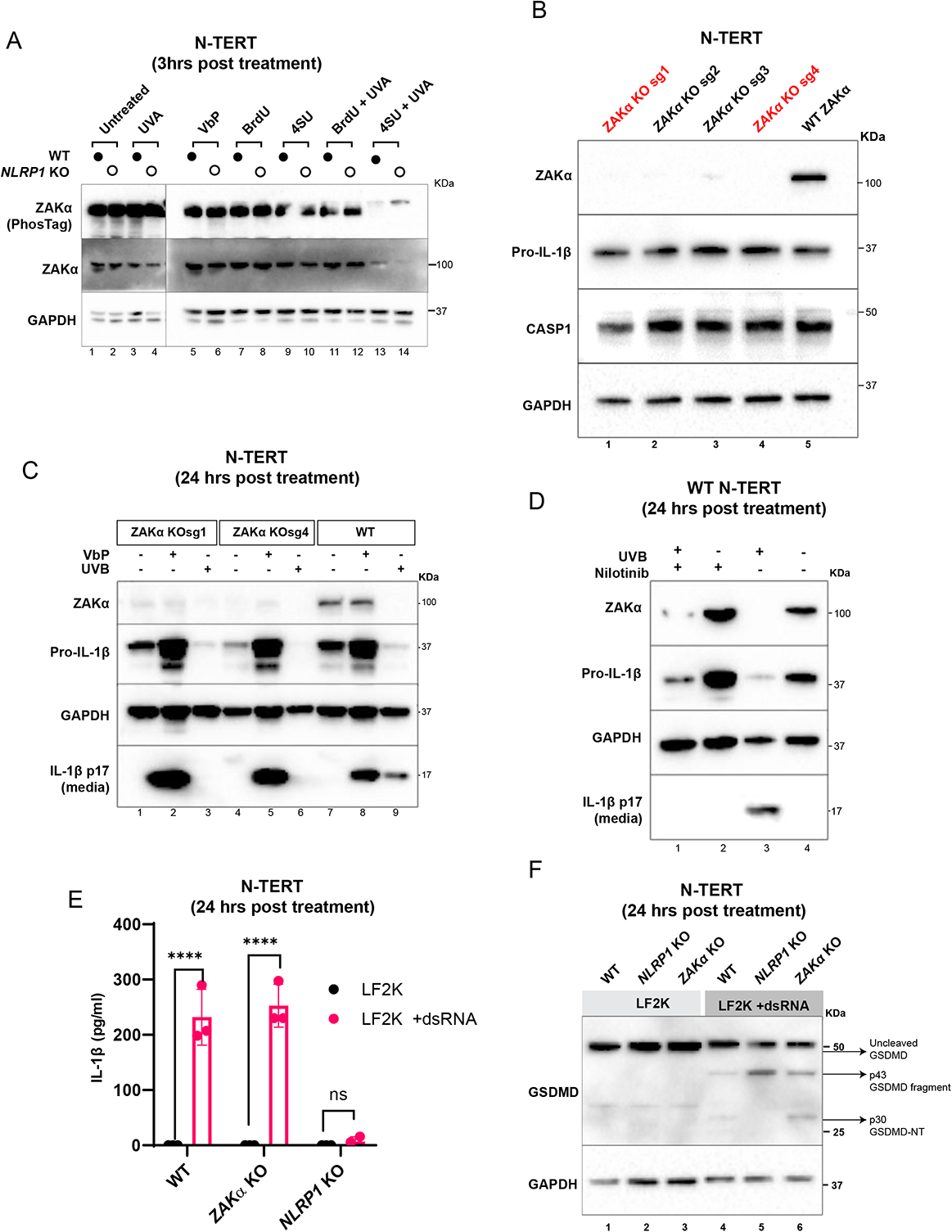
Additional evidence that RSR kinase ZAKɑ controls UVB-but not VbP- or dsRNA-triggered NLRP1 activation. A. Immunoblot of the same lysates used in the selective photosensitization experiment described in Figure 2C. B. Immunoblot validation of WT or *ZAKɑ* KO N-TERT cells. Red indicates *ZAKɑ* KO cells that were used for further study. C. Immunoblot of WT or *ZAKɑ* KO N-TERT treated with UVB (100mJ/cm^2^), VbP (2 μM) or sham irradiated and harvested 24 hours later. D. Immunoblot of WT or *ZAKɑ* KO N-TERT treated with UVB (100mJ/cm^2^) with or without Nilotinib (0.1μM), or sham irradiated. Cells were harvested 24 hours later. E. IL-1β ELISA of WT, *ZAKɑ* KO or *NLRP1* KO N-TERT treated with lipofectamine alone (LF2K) or LF2K complexed with dsRNA (LF2K +dsRNA) and harvested 24 hours later. F. Immunoblot of cell lysates from E.

**Figure S4.**
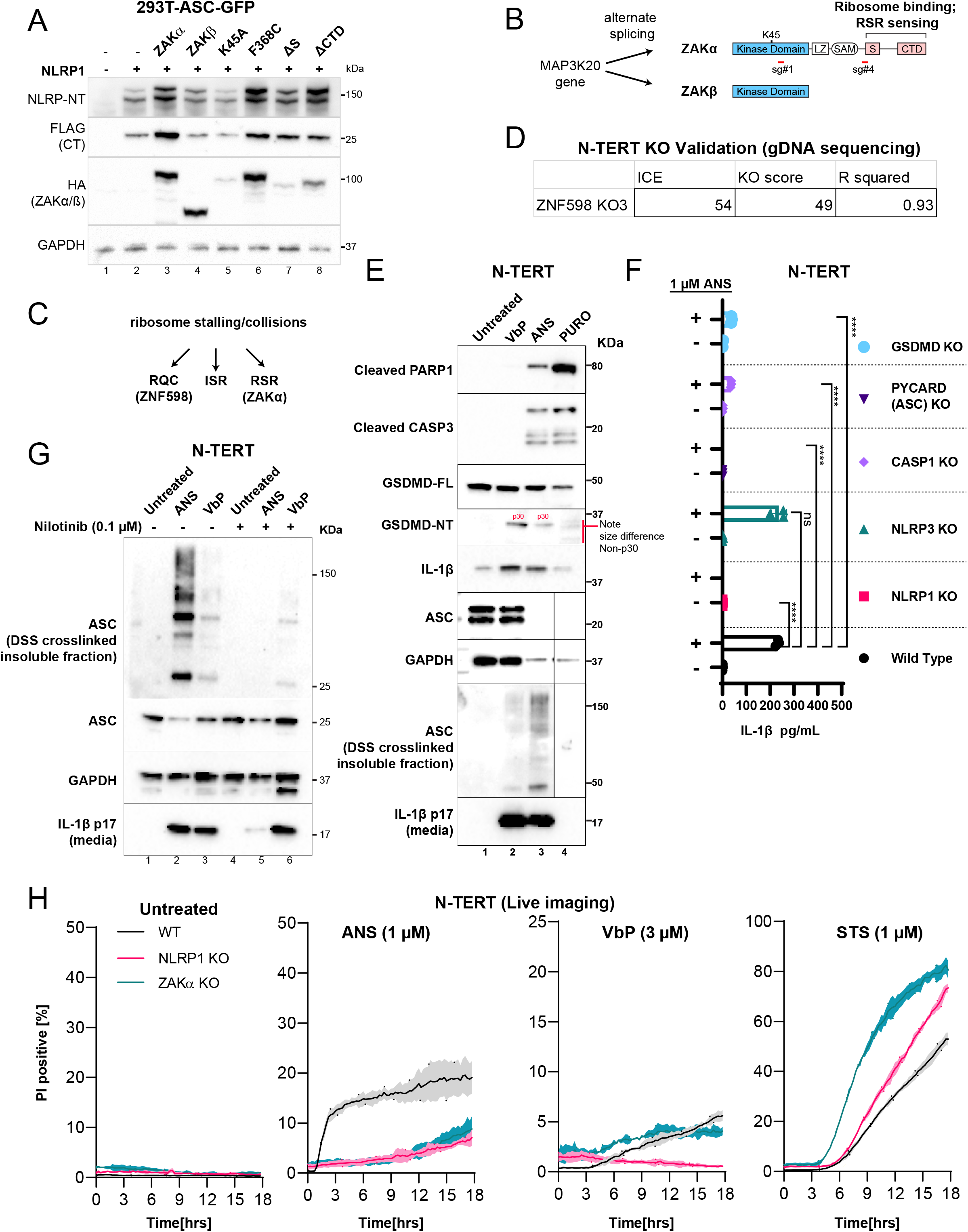
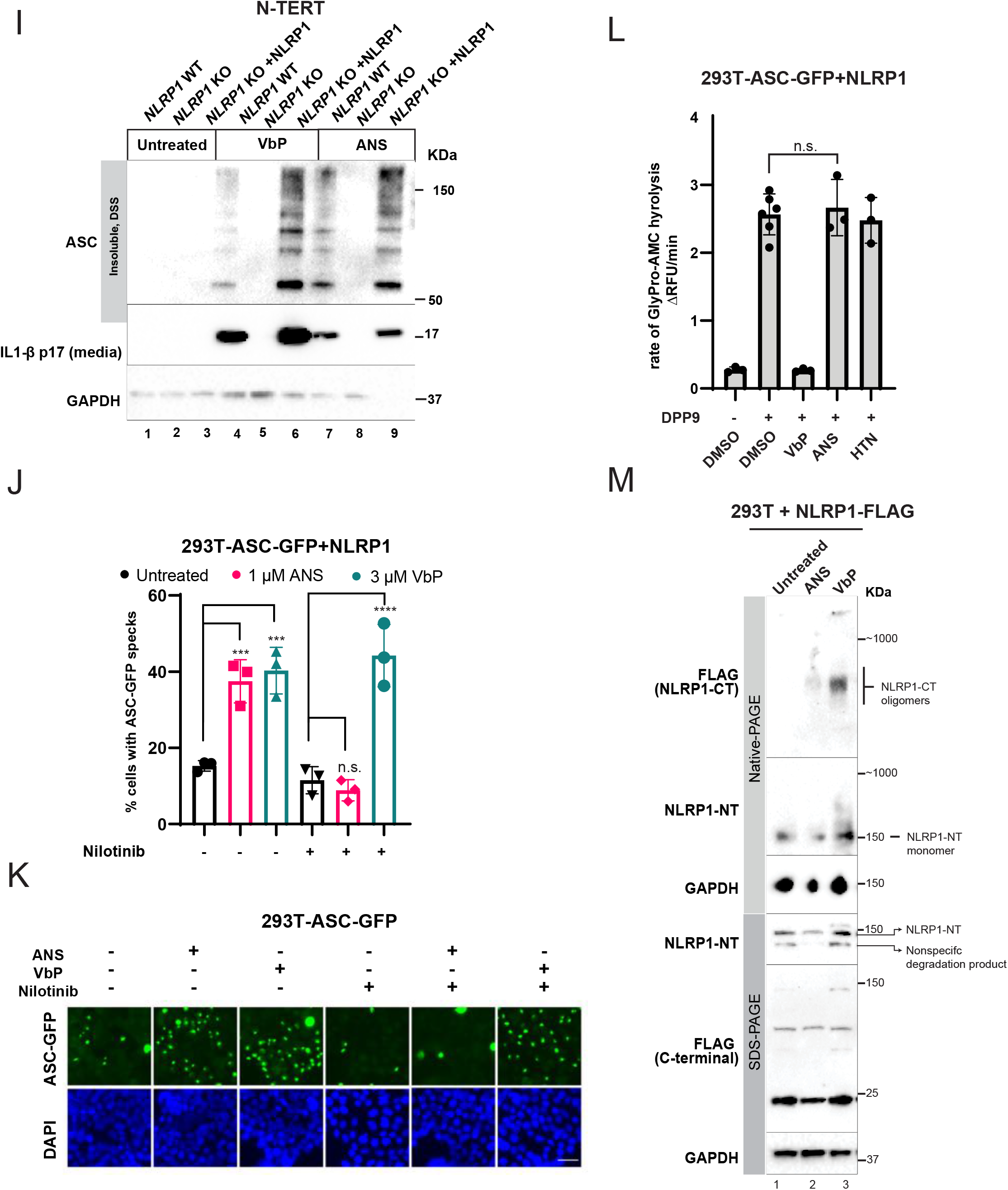
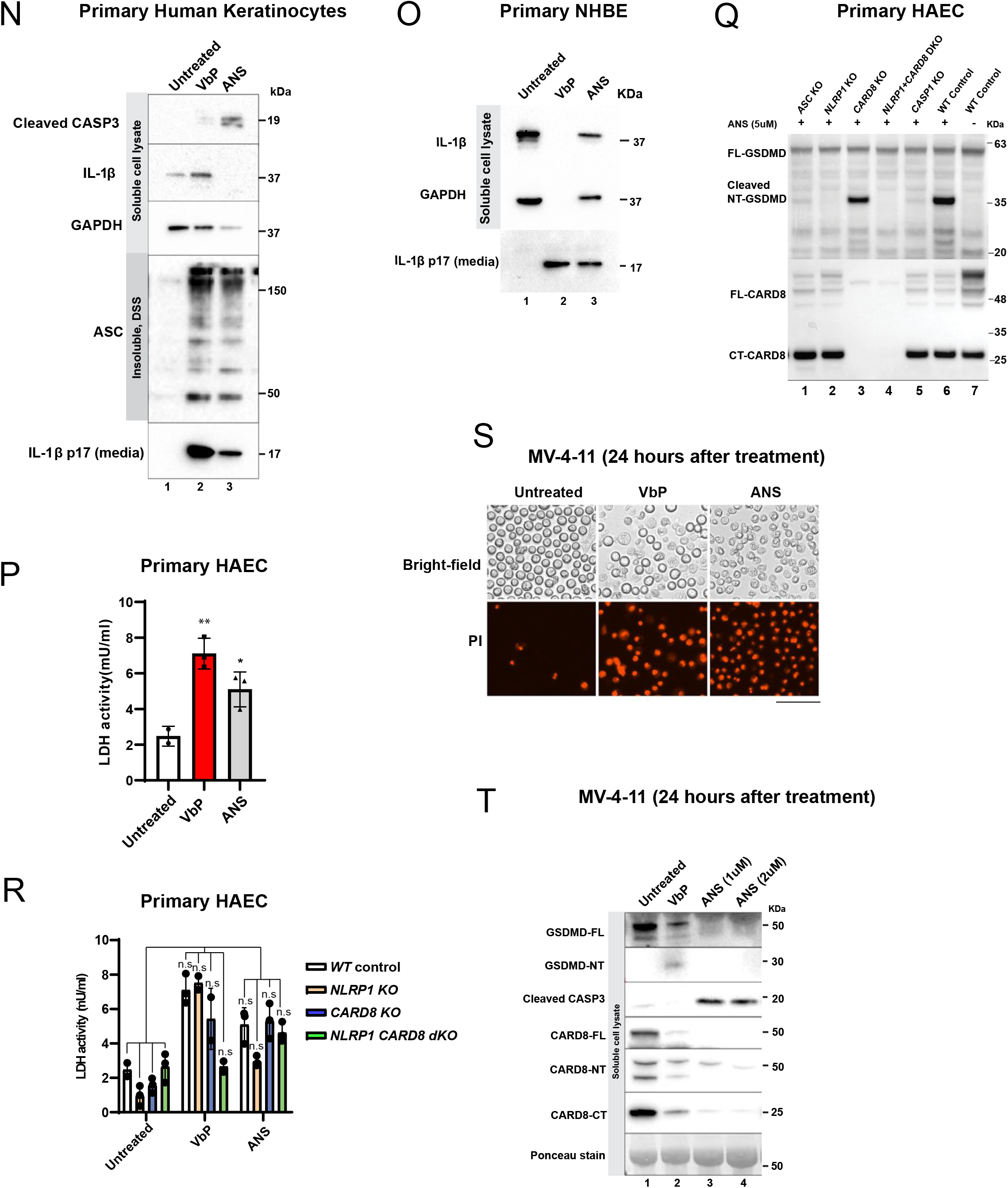
Additional evidence demonstrating that over-expression and chemical activation of ZAKɑ triggers the NLRP1 inflammasome. A. Immunoblot from 293T-ASC-GFP cells transfected with the indicated plasmids. Cells were harvested 48 hours post transfection. B. The target sites of ZAKɑ CRISPR guide RNAs in relation to ZAKɑ and ZAKβ domain structures. C. Summary of the 3 main pathways induced by ribosome stalling and collisions and their key regulators D. Knockout validation of the *ZNF598* KO N-TERT via Sanger sequencing of the genomic DNA using the Synthego ICE tool. E. Immunoblot from ANS-, VbP-, and PURO-treated N-TERT lysate. DSS (1 mM) crosslinking was performed on NP40 insoluble pellets after lysis. Note that PURO caused non-p30 GSDMD cleavage. FOr ANS and GAPDH, samples were run on the sample blot (instead on spliced from different blots) with irrelevant lanes cropped out. F. IL-1β ELISA from untreated or ANS-treated N-TERT cells of the indicated genotypes. G. Immunoblot from lysates, crosslinked pellets and culture media of wild-type N-TERT cells treated with the indicated drugs. Nilotinib pretreatment was initiated 30 mins before the addition of ANS and VbP. H. Quantification of PI positive N-TERT cells treated with the indicated drugs. Cells were imaged at 15 min intervals for 18 hours. I. Immunoblot from lysates, crosslinked pellets and culture media of wild-type, *NLRP1* KO and NLRP1-expressing *NLRP1* KO N-TERT cells treated with the indicated drugs. J. Percentage of ASC-GFP speck forming cells in 293T-ASC-GFP cells transfected with NLRP1 and treated with the indicated drugs. K. Representative images of J. L. Cell-free DPP9 activity assay measured by the rate of GlyPro-AMC hydrolysis, from 293T-ASC-GFP-NLRP1 lysates incubated with the indicated drugs. HTN=harringtonin, 1 μM. M. Immunoblot following SDS-PAGE or Native-PAGE of 293T-NLRP1-FLAG lysates treated with the indicated drugs. Cells were harvested 5 hours post drug treatment. N. Immunoblot from soluble lysate, insoluble pellets and culture media of primary human keratinocytes treated with 3 μM VbP or 1 μM ANS for 24 hours. O. Immunoblot from soluble lysate, insoluble pellets and culture media of primary human bronchial epithelial cells (NHBEs) treated with 3 μM VbP or 1 μM ANS for 24 hours. P. LDH activity in the culture media of primary human aortic endothelial cells (HAECs) treated with 3 μM VbP or 1 μM ANS for 24 hours. Q. Immunoblot from soluble lysate of various CRISPR/Cas9 KO human aortic endothelial cells (HAECs) treated with VbP or ANS. R. LDH activity in the culture media of HAECs shown in Q. S. PI inclusion of MV-4-11 cells treated with VbP and ANS. T. Immunoblot of MV-4-11 cells treated with VbP and ANS as shown in S.

**Figure S5.**
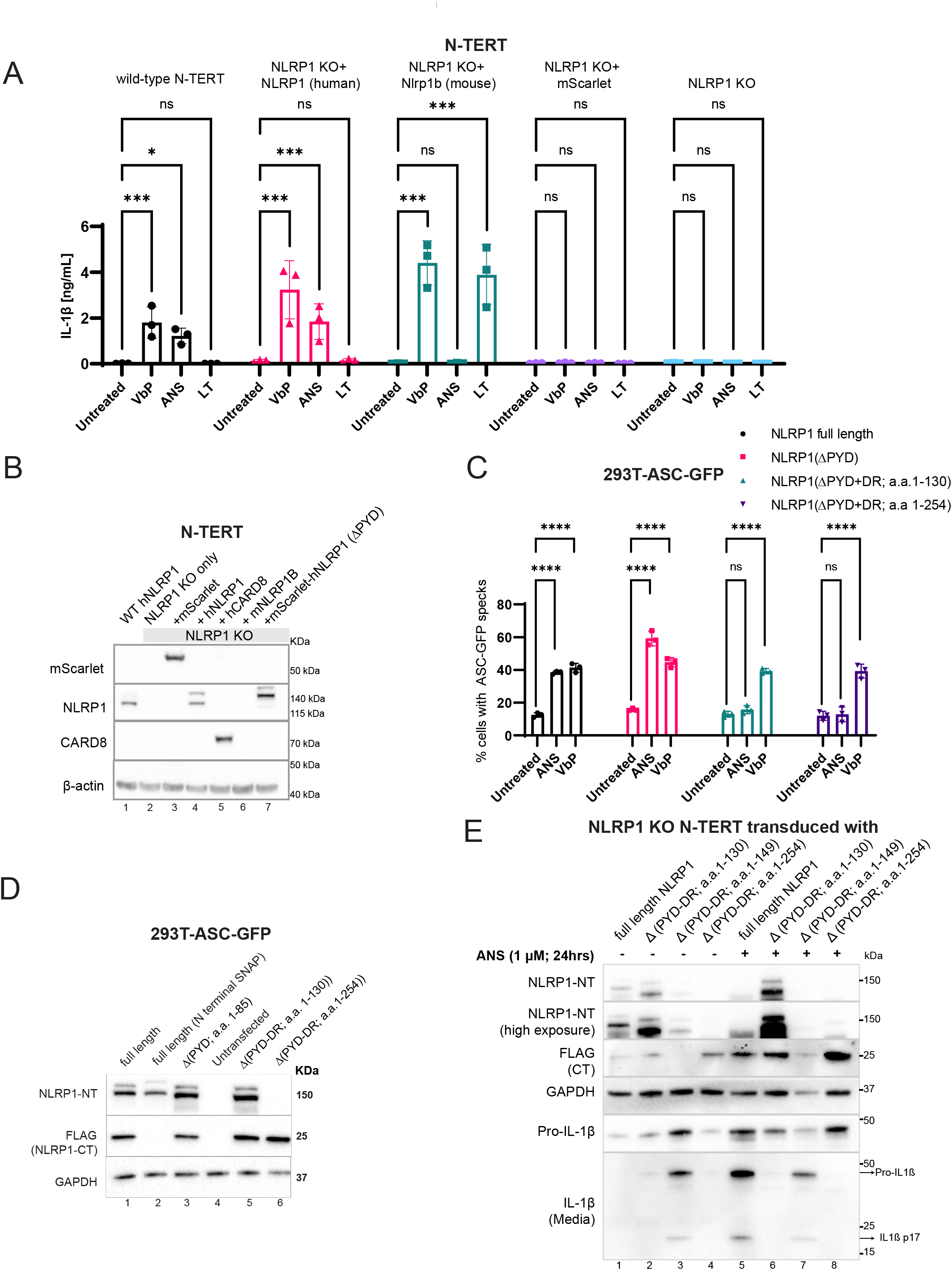
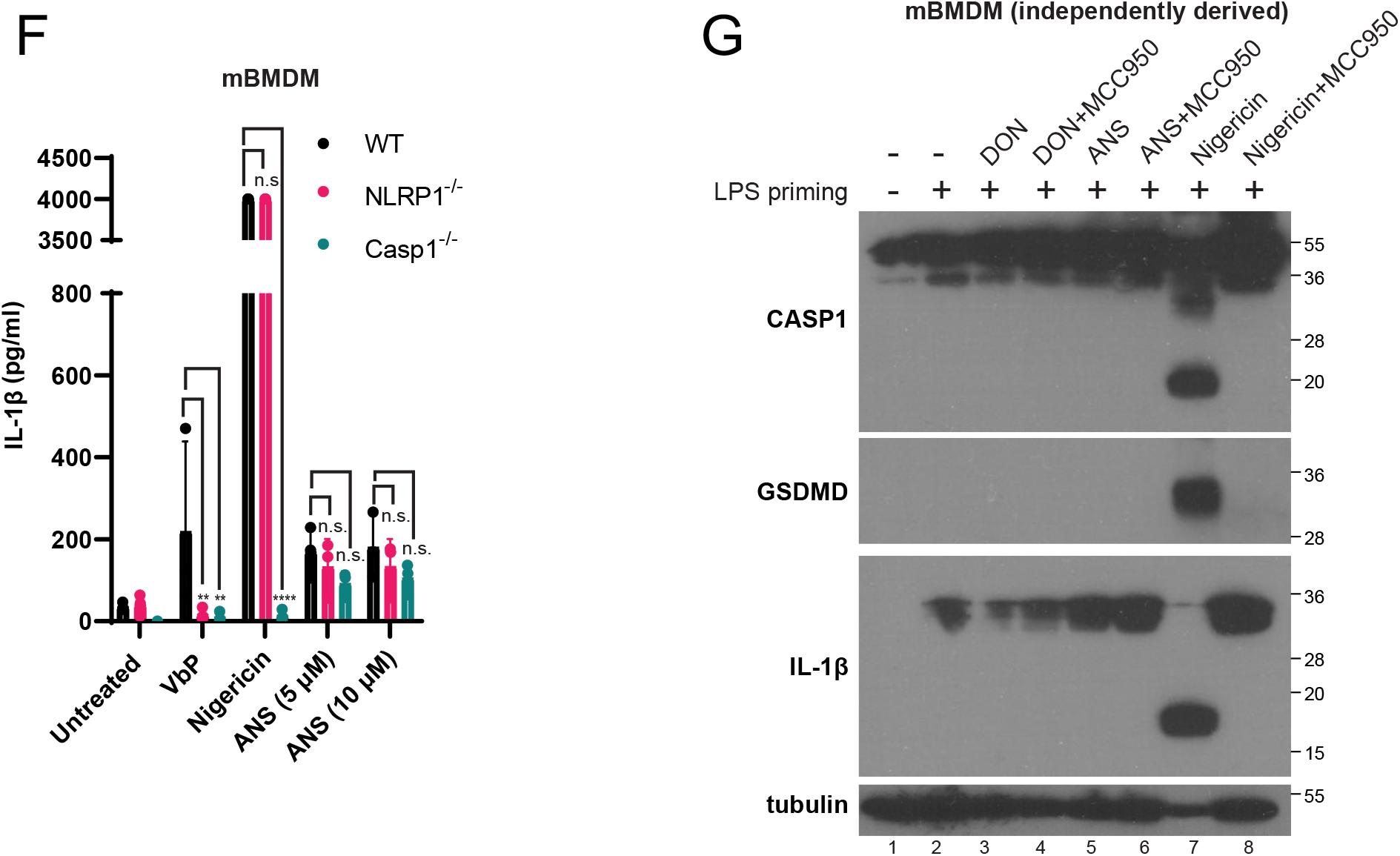
Additional evidence supporting the requirement of NLRP1^DR^ in ZAKɑ-dependent, but not VbP-dependent NLRP1 activation. A. IL-1β ELISA from *NLRP1* KO N-TERT cells reconstituted with the indicated NLRP1 variants and treated with LF. B. Immunoblot from cells shown in Figure 5A and S5A. C. Percentage of ASC-GFP speck forming cells in 293T-ASC-GFP cells transfected with the indicated NLRP1 mutants and treated with ANS or VbP. D. Immunoblot of 293T-ASC-GFP cells shown in C. Note that the epitopes of the NLRP1-NT antibody resides in NLRP1^DR^ (most likely a.a. 130-254). Hence it does not effectively detect the Δ(PYD-DR, a.a. 1-254) mutant shown in lane 6. This should not be interpreted as a loss of protein accumulation. E. Immunoblot of lysate made from independently generated *NLRP1* KO N-TERT cells expressing different DR mutants. Note that these constructs have the C-terminal FLAG tag but not the N-terminal GFP tag used in Figure 5. As a result, the ΔPYD-DR (a.a. 1-254) mutant could not be detected effectively by the NLRP1 NT antibody whose major epitopes reside in this region. This should not be interpreted as a loss of protein accumulation. In fact, both ΔPYD-DR mutants show increased abundance of NLRP1-NT after ANS treatment in contrast for wild-type NLRP1 (lanes 2 vs. 6, lane 4 vs 8, NLRP1 NT panel). The reason for this stabilization is not clear currently. F. Murine IL-1β ELISA from Nlrp1a-c^-/-^ and Casp1^-/-^ BMDMs treated with the indicated drugs. G. Immunoblot of mixed supernatant and extracts. Independently generated C57BL/6 BMDMs were primed with 100 ng/mL LPS for 3 hours and stimulated with the indicated drugs in the presence or absence of MCC950.

**Figure S6.**
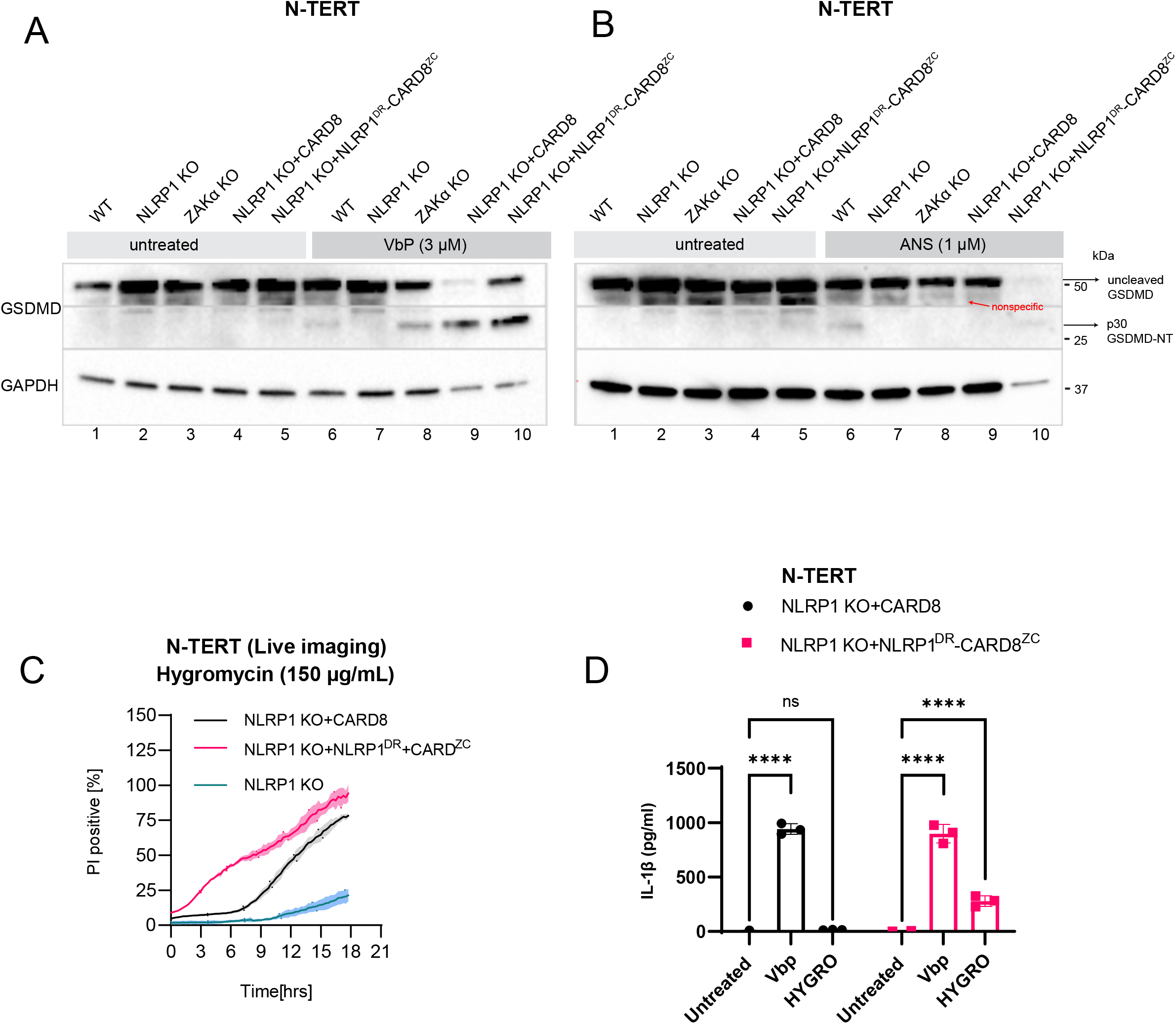
Further characterization of NLRP1^DR^-CARD8^ZC^. A. GSDMD immunoblot from untreated and VbP-treated N-TERT lysates of the indicated genotypes. Cells were harvested 24 hours after treatment. The p30 fragment is marked with black arrow. B. GSDMD immunoblot from untreated and ANS-treated N-TERT lysates. C. Quantification of PI positive cells in N-TERT cells expressing the indicated sensors during the 18 hour incubation with hygromycin. D. IL-1β ELISA from VbP- or HYGRO-treated N-TERT cells of the indicated genotypes 24 hours post treatment.

**Figure S7.**
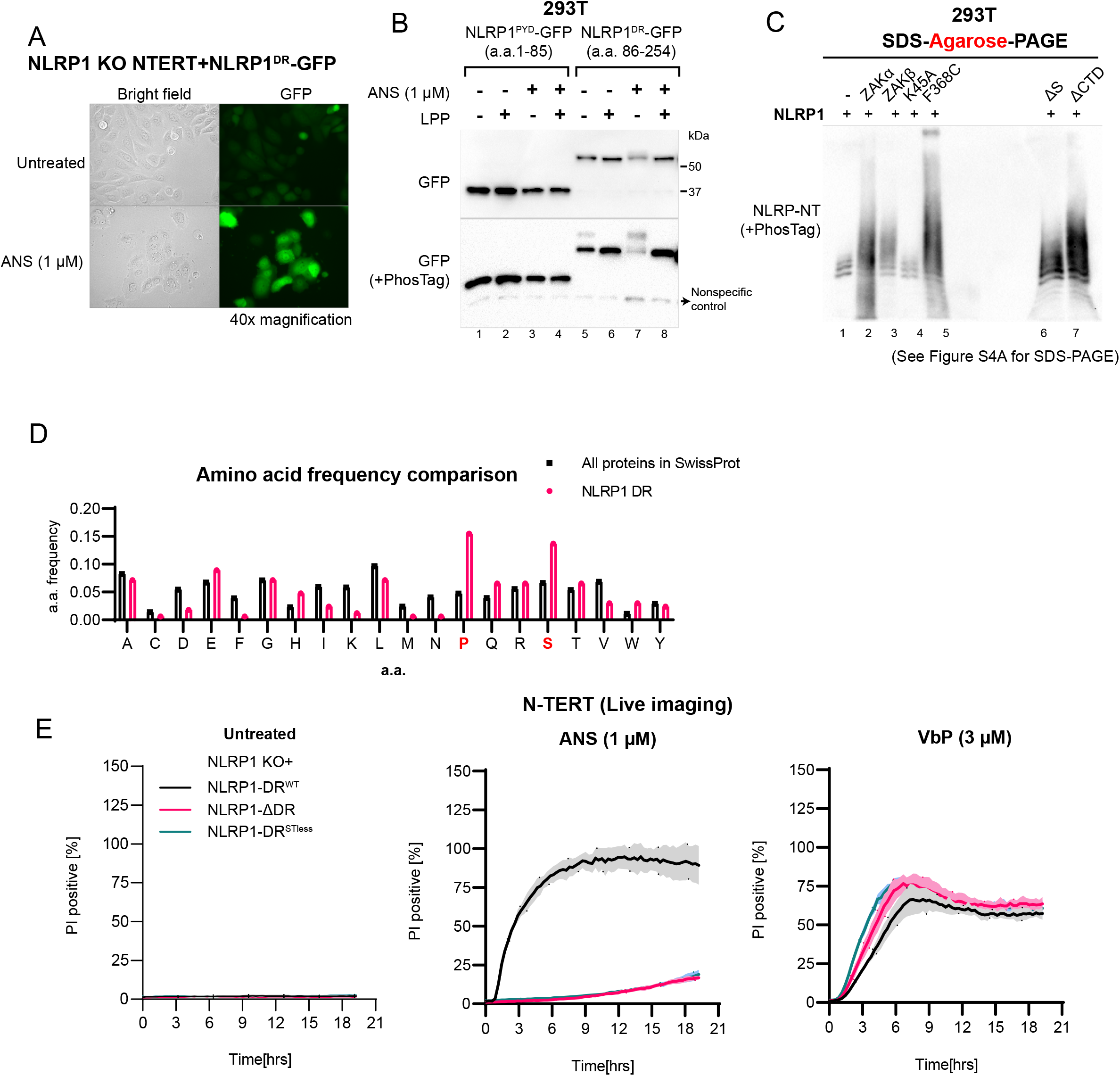
Additional evidence on the requirement of NLRP1^DR^ hyperphosphorylation in ZAKɑ-dependent NLRP1 activation. A. GFP fluorescence and bright field images of *NLRP1* KO N-TERT cells expressing NLRP1^DR^ (a.a. 86-254)-GFP before and after ANS treatment (1 μM) for 24 hours. GFP fluorescence images were acquired using the same exposure time. B. GFP Immunoblot following conventional SDS-PAGE or PhosTag SDS-PAGE of 293T lysates expressing the indicated GFP fusion constructs. Cells were harvested 3 hours post ANS treatment. Lambda phosphatase (LPP) was added post-lysis for 30 mins. C. NLRP1-NT immunoblot following PhosTag containing SDS-Agarose-PAGE. Lysates were prepared from the experiment shown in Figure 4B. D. Comparison of the individual amino acid frequencies of NLRP1^DR^ (a.a. 86-254) relative to all proteins in SwissProt. E. Quantification of the percentage of PI positive *NLRP1* KO cells expressing the indicated set of NLRP1 ΔPYD DR mutants. Images were acquired at 15 min intervals for 18 hours in the presence of ANS, VbP and PI.

